# Rapid folding of nascent RNA regulates eukaryotic RNA biogenesis

**DOI:** 10.1101/2024.11.26.625435

**Authors:** Leonard Schärfen, Isaac W. Vock, Matthew D. Simon, Karla M. Neugebauer

## Abstract

An RNA’s catalytic, regulatory, or coding potential depends on RNA structure formation. Because base pairing occurs during transcription, early structural states can govern RNA processing events and dictate the formation of functional conformations. These co-transcriptional states remain unknown. Here, we develop CoSTseq, which detects nascent RNA base pairing within and upon exit from RNA polymerases (Pols) transcriptome-wide in living yeast cells. By monitoring each nucleotide’s base pairing activity during transcription, we identify distinct classes of behaviors. While 47% of rRNA nucleotides remain unpaired, rapid and delayed base pairing – with rates of 48.5 and 13.2 kb^-1^ of transcribed rDNA, respectively – typically completes when Pol I is only 25 bp downstream. We show that helicases act immediately to remodel structures across the rDNA locus and facilitate ribosome biogenesis. In contrast, nascent pre-mRNAs attain local structures indistinguishable from mature mRNAs, suggesting that refolding behind elongating ribosomes resembles co-transcriptional folding behind Pol II.

## Introduction

Eukaryotic transcription occurs in the nucleus and is catalyzed by three different RNA polymerase holoenzymes (Pols I, II and III) that are nevertheless similar in structure and function.^1^ All classes of RNA are extensively processed starting during transcription, resulting in rapid covalent modification of the molecule or editing of its sequence. In addition to covalent editing, intramolecular base pairing, which drives RNA folding, is a chemical property of RNA that is key to its cellular function.^2,3^ In mRNA, base pairing can extend RNA half-lives and/or regulate translation, while single-strandedness may be necessary to enable access to processing signals where recognition is sequence based.^4,5^ Highly folded RNAs like rRNA, snRNAs and tRNAs require base-pairing to form structures that associate with proteins or achieve catalytic activity. Due to limitations in detecting dynamic RNA base pairing during synthesis and specifically in cells, how and when RNA structures first arise and later mature, and how proteins remodel them to facilitate processing steps is currently unknown. Several examples of potentially disease-causing disruptions in gene expression due to misfolding were recently described^6,7^, but we lack a basic understanding of the earliest folding events: prevalence, timing, and persistence of nascent RNA structures in eukaryotes.

Co-transcriptional base pairing has the potential to regulate all eukaryotic RNA processing steps that begin and are often completed during transcription, including pre-mRNA splicing, RNA editing and modification, and 3’-end cleavage.^8,9^ Recent work has established that base pairing is often dynamic in living cells, and RNAs can populate multiple structural states while sharing identical sequence.^10–13^ Consequently, prediction of base pairing based on sequence alone often fails to reflect true structures due to the rough energy landscape of RNA folding as well as extensive RNA processing. Thus, recent efforts have focused on chemically probing RNA structure *in vitro* and *in vivo*. Dimethyl sulfate (DMS) or SHAPE (selective 2′-hydroxyl acylation analyzed by primer extension), which modify unpaired nucleobases at the Watson–Crick face or at the 2′-hydroxyl backbone position, respectively, are broadly utilized to measure base pairing *in vivo*. These methods have mostly been applied to mature, fully folded RNA – but understanding how base pairs initially arise is equally critical for harnessing the full potential of eukaryotic RNA for therapeutic or engineering purposes, and will enable their manipulation.^14^

The process of RNA folding should begin co-transcriptionally, given faster timescales of base pairing compared to RNA synthesis.^15^ To date, nascent RNA base pairing has mostly been studied in bacteria where co-transcriptional regulation of transcription and translation is well established.^16–18^ For example, pioneering studies in prokaryotes have revealed that local RNA folding during transcription can to promote or antagonize mature RNA structure due to transient or more permanent population of local energy minima prior to full-length synthesis.^19–21^ Studies in eukaryotes have been focused on individual examples, and indeed, changes in transcription speed affect eukaryotic stem-loop formation important for 3’end cleavage of nascent histone pre-mRNAs.^22^ This singular case highlights the currently underappreciated regulatory power of co-transcriptional base pairing.

Experimental approaches to studying folding in eukaryotic rather than prokaryotic systems must contend with differences in the genomes and regulatory mechanisms present. The scarcity of eukaryotic nascent RNA has precluded high-resolution *a priori* analyses of nascent RNA structure, especially in living cells. Moreover, eukaryotic genomes are larger, gene architectures are more complicated, and multiple RNA polymerases operate with distinct regulatory machineries.^9^ Because of these challenges, previous approaches have included probing nuclear RNA^23^ or isolating nascent RNA and then re-folding and probing it in vitro^22^; these methods treat nascent RNA as a pool and thereby lose the connection of base pairing to polymerase position. In contrast, correlation of RNA polymerase density with computationally predicted structure has suggested a connection between folding and elongation rates.^24^ Thus, the lack of methods to directly observe RNA base pairing during transcription weakens interpretations of current data. RNA synthesis within a highly organized and crowded nucleus likely differs from these experimental conditions, since many co-transcriptional regulatory mechanisms are features of endogenous gene loci packaged in cellular chromatin. Here, we overcome these obstacles by establishing a new method, Co-transcriptional Structure Tracking (CoSTseq) that measures base pairing as a function of the position of the three eukaryotic RNA Pols, enabling an analysis of co-transcriptional regulatory events related to RNA structure in living cells. Additionally, CoSTseq’s unprecedented accuracy of methylation rate profiling within RNA Pol holoenzymes identifies the key structural features of the RNA’s path through the transcription bubble and exit channel, where we find that folding is unfavorable. Using CoSTseq, we test the functions of factors widely hypothesized to contribute to intramolecular RNA folding: transcription elongation factors and RNA helicases.^25–27^

## Results

### Co-transcriptional Structure Tracking (CoSTseq) detects nascent RNA base pairing

To understand nascent RNA base pairing in eukaryotes, we devised a biochemical strategy based on biotin-NTP transcriptional run-on and dimethyl sulfate (DMS) treatment (Fig. 1a). CoSTseq enables isolation of nascent RNA molecules that are methylated at unpaired residues, in addition to identification of the position of RNA Pols by a biotinylated nucleotide. To achieve this, *Saccharomyces cerevisiae* cells were permeabilized and incubated with biotinylated CTP such that incorporation of the biotinylated nucleotide marks the nascent RNA 3’-end.^28^ The cells were then treated with DMS to methylate unpaired residues. Biotinylated nascent RNA was affinity purified from total RNA and reverse transcribed using GsI2c reverse transcriptase (RT) from *Geobacillus stearothermophilus*^29^ using its template switching ability and re-coding of methylation events at A, C and U residues into mutations in cDNA. A 5’-end adapter with 7xN unique molecular identifier (UMI) was ligated to the cDNA, which was then read by high-throughput sequencing (Fig. 1a). The 3’-end distribution across genes combined with RT fall-off yields cDNA fragments of optimal length for Illumina 2×150 paired-end sequencing, obviating the need for fragmentation (Fig. S1a). For every CoSTseq dataset we also performed DMS-MaPseq in parallel to probe mature RNA structure^30^ and compare it to nascent RNA. Through dual encoding of RNA Pol position and base pairing status, each CoSTseq read contains both types of information. While every read can be directly assigned to a Pol position, interpretation of base pairing status requires sufficient sequencing coverage to calculate mutation rates (see Methods). Analogous to DMS-MaPseq^30^, mutation rates were calculated by dividing the number of mutations by the sequencing depth at each position. Raw mutation rates for positions with at least 900 aligned reads for A, C and 2,000 for U nucleotides were normalized, yielding DMS reactivity values that range from 0 (likely paired) to >1 (likely unpaired).

**Fig. 1:**
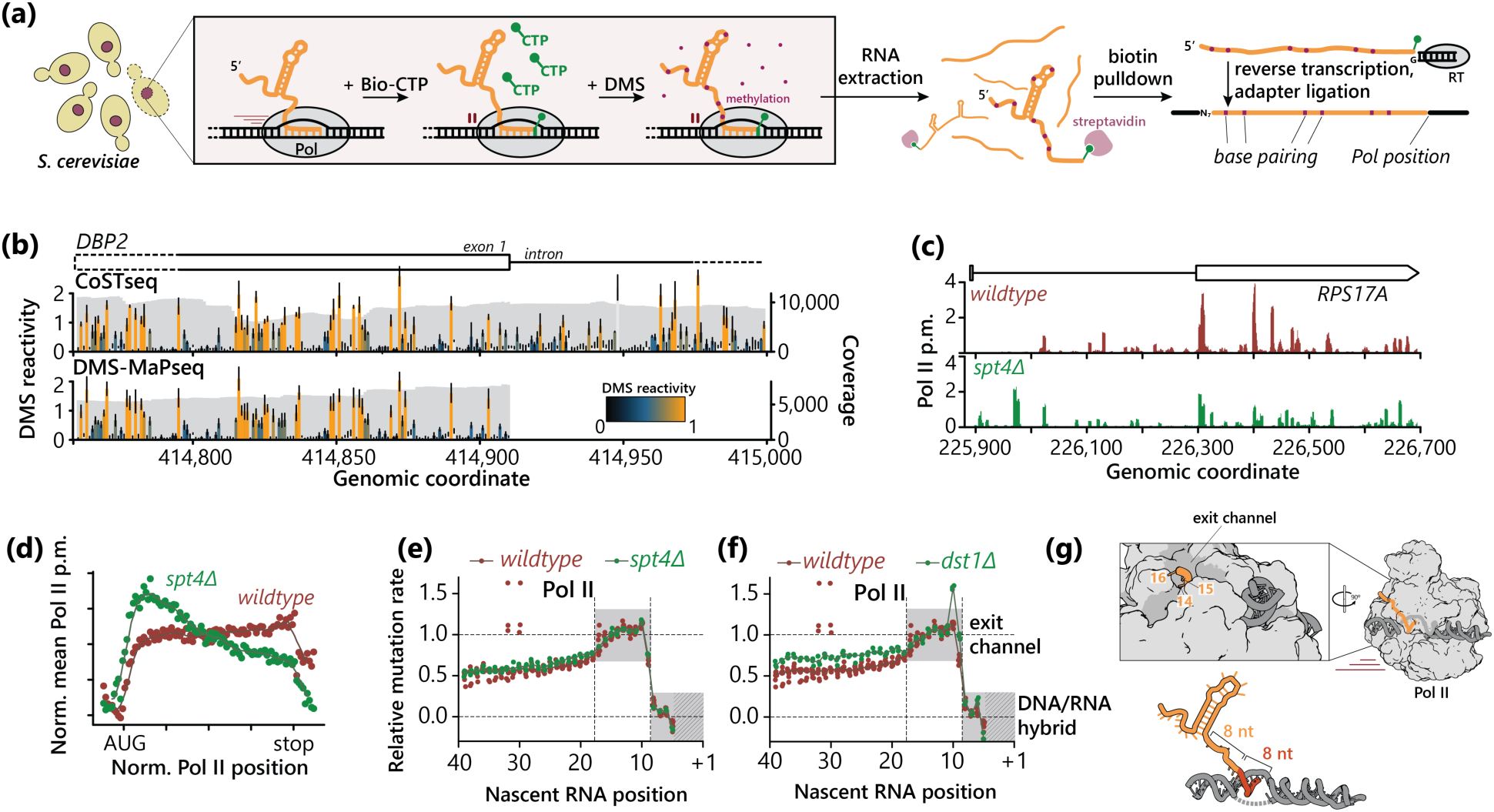
CoSTseq detects local RNA base pairing, which begins 16 nucleotides from the catalytic center of Pol II. **(a)** Yeast cells are permeabilized and treated with biotinylated CTP, then DMS. RNA is extracted and then enriched with streptavidin. Nascent RNA is reverse transcribed, a 5’-end adapter is ligated, and the resulting cDNA is amplified for Illumina sequencing. **(b)** DMS reactivity profiles from CoSTseq (nascent RNA, projected across RNA Pol II positions) and DMS-MaPseq (mature RNA) reads. Light gray bars indicate G/U. Bars at A/C nucleotides are colored according to normalized signal. Error bars represent the standard deviation from at least three biological replicates at each position. CoSTseq: n=8, DMS-MaPseq: n=3. Read coverage is plotted in dark gray. **(c)** 3’-end (RNA Pol II) positions from CoSTseq reads for wildtype and *spt4Δ* strains. 3’-end coverage is normalized according to total sequencing depth (reported as per million, p.m.) and summed over a six-nucleotide sliding window. **(d)** 3’-end meta gene profiles for 100 bins per transcript between start and stop codon. **(e)** Mutation rates for nucleotides close to the RNA 3’-end (Pol II active site, +1) calculated from all reads aligned to Pol II transcribed genes. Mutation rate signal is normalized between the mean of positions 6-8 (set to zero) and 12-18 (set to one). Dots indicate individual biological replicates; solid line represents the median. Wildtype: n=8, *spt4Δ*: n=3. **(f)** Wildtype: n=8, *dst1Δ*: n=3. **(g)** Cartoon based on the molecular structure of the Pol II holoenzyme^36^, showing the path of the nascent RNA from the catalytic center to the RNA/DNA hybrid to the exit channel, where nucleotide 16 is the first to reach the surface.

To visualize the quality of the obtained data we began by validating our ability to monitor Pol II position and mutation rate in nascent pre-mRNA. Projected CoSTseq DMS reactivity profiles are generated by summing CoSTseq signal (number of mutations, coverage) across RNA Pol positions, yielding a profile that represents average base pairing information for nascent RNA across the region. These profiles are analogous to what DMS-MaPseq reports for mature RNA, and projected CoSTseq DMS reactivity profiles show similar signal-to-noise ratio (Fig. S1b). We are confident that CoSTseq reports on RNA Pol II activity because these profiles capture nascent RNA base pairing within introns, which are quickly and co-transcriptionally removed from nascent RNA in yeast^31^ (Fig. 1b). To analyze elongation rate and provide a quality control for our biotinylation strategy, we performed CoSTseq and DMS-MaPseq in a fresh *spt4Δ* knockout strain and mapped coordinate of the 3’-end nucleotide for every aligned read. Spt4 is an RNA Pol II elongation factor, and its absence leads to a characteristic change in the slope of RNA Pol II density across gene bodies relative to wildtype.^32^ This phenotype is evident from both the CoSTseq 3’-end profiles for individual genes (Fig. 1c), as well as the metagene profile that shows a steeper negative slope for RNA Pol II density along the length of genes, as expected for slower elongation rates^28^ (Fig. 1d). We conclude that CoSTseq can capture RNA Pol positions at C nucleotides with high enough accuracy to detect changes in transcriptional kinetics and with high reproducibility (average Pearson’s correlation coefficient (PCC) for biological replicates of 0.94).

### Intramolecular nascent RNA base pairing begins upon exit from Pols I and II

Previous studies have indicated diminished reactivity of nascent RNA when base paired with DNA in the transcription bubble, as expected.^16,17,33,34^ Owing to the high precision and coverage of RNA Pol catalytic sites obtained by CoSTseq, we sought to obtain insight into the path of nascent RNA within eukaryotic RNA polymerases and to determine the position of Pols I and II when nascent RNA folds for the first time. To analyze the reactivity of nascent RNA within RNA Pol II, reads that align to genes transcribed by RNA Pol II are piled up at their 3’-ends, yielding a population-level DMS signal representing a footprint relative to the holoenzyme’s active site (Fig. 1e). We find an eight-nucleotide region of initially low reactivity, which increases to high-reactivity for eight nucleotides and then progressively drops off to an intermediate level. These distinct footprints correspond to structural models of transcribing RNA Pol II, with eight nucleotides protected in the DNA/RNA hybrid and 6-8 nucleotides in the exit channel.^35,36^ These data imply that DMS enters the exit channel, in which base pairing is physically constrained. The drop in reactivity between nucleotides 16-20 indicates that each nucleotide is available for base pairing at the moment of its exit from RNA Pol II, and that base pairing becomes more likely with increasing distance to the exit channel. Importantly, we do not observe a similar reactivity decrease from DMS-MaPseq reads that are randomly fragmented and do not carry 3’-end information, or from untreated samples (Fig. S1c), showing that these footprints are specific for nascent RNA as it travels through elongating RNA Pol holoenzymes. The similarity between the RNA Pol I and II CoSTseq reactivity footprints (Fig. S1d) is consistent with similar exit channel paths for nascent pre-rRNA and pre-mRNA through RNA Pols I and II, respectively.

In prokaryotes, transcriptional pausing promotes local structure formation (see Introduction), which may take place within the exit channel of bacterial polymerase.^37^ To address the possibility that folding might take place within eukaryotic RNA Pol II if given more time, we analyzed CoSTseq data from the *spt4Δ* deletion mutant and found it has no effect on the pattern of DMS reactivity either within the exit channel or after exit from RNA Pols I and II (Fig. 1e, Fig. S1d). We also created a fresh strain lacking the elongation factor TFIIS (*dst1Δ*) and generated CoSTseq and matching DMS-MaPseq datasets in three biological replicates. TFIIS is a Pol II elongation factor that recovers Pol II from backtracked, paused states by cleaving the nascent RNA inside the polymerase.^35,38^ Unexpectedly, CoSTseq reveals an altered Pol II footprint at position 10 of Pol II (Fig. 1f), suggesting greater availability of this nucleotide in pre-mRNA when TFIIS is absent. This change may be due to altered contacts with the Pol II Rbp1 subunit and/or decreased base pairing with DNA as it leaves the RNA/DNA helix within the transcription bubble. Thus, CoSTseq data agrees with and extends RNA Pol I and II atomic structures and molecular mechanisms (Fig. 1g), highlighting their ability to accurately reveal co-transcriptional folding and novel aspects of the RNA’s path.

### Co-transcriptionally formed long-range interactions define the structure of yeast U2 snRNA

Given the potential for rapid folding, we were intrigued by the question of how base pairs that require long-range interactions form co-transcriptionally. *S. cerevisiae* U2 snRNA (*lsr1* gene) is transcribed by Pol II and uniquely carries a nearly 1 kb insertion, which is flanked by the conserved functional structure elements.^39,40^ This insertion is not required for U2 snRNA function^41^, but its size suggests that it must form a distinct structure or be otherwise sequestered to avoid a disruption of the functional folding pattern of U2. To monitor how base pairs arise in U2, we calculated average DMS reactivity and Gini index across the U2 locus. Low Gini index (evenly distributed reactivity values) and high DMS reactivity suggest flexible, unstructured regions while high Gini index and low DMS reactivity suggest highly defined, base paired regions. CoSTseq data support a model whereby the 5’-half of the insert remains unpaired during transcription, while the second half base pairs immediately (Fig. 2a): in the first half of the gene, Gini index and average DMS reactivity are initially low and high, respectively. Once the turning point is reached, the Gini index increases and DMS reactivity falls off, consistent with co-transcriptional base pairing and stem formation. Additionally, the first half is particularly U-rich while the second half is enriched in U-complementary A and G residues (Fig. 2b), consistent with base pairing between the two halves. Indeed, structure prediction based on DMS reactivities of only the second half of the insert reveal a large 440 base pair (927 nt) stem capped with a distinct 3-stem junction (Fig. 2c). Co-transcriptional sequestration of the insert sequence through base pairing likely protects the downstream functional part of U2 snRNA from engaging in off-target pairings during transcription, processing and RNP assembly (Fig. 2d). This structure might also prevent off-pathway interactions during the splicing cycle.

**Fig. 2:**
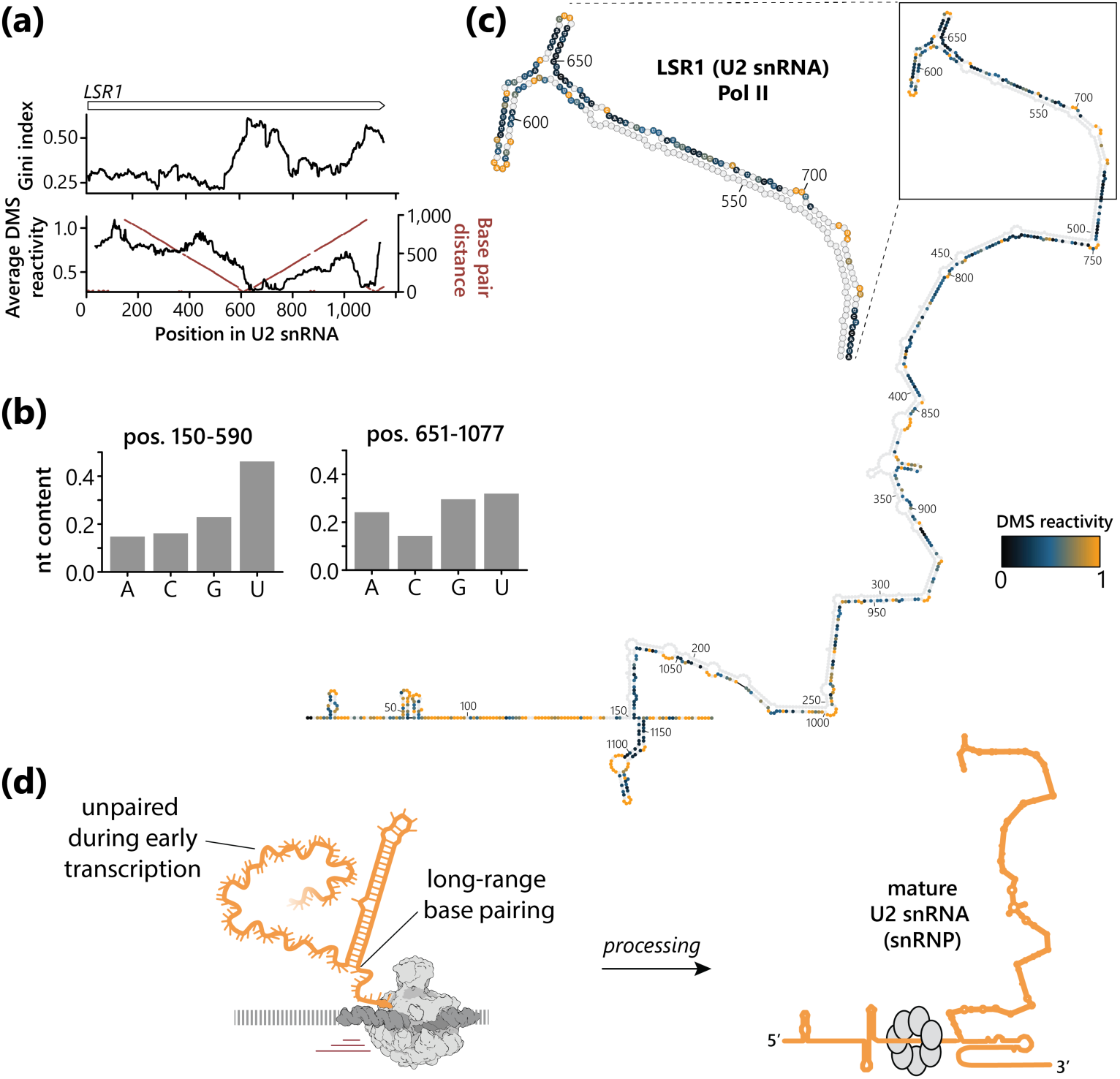
Co-transcriptional formation of a long-range stem loop in U2 snRNA *in vivo*. **(a)** Gini index and average DMS reactivity in windows of size 70 across the nascent U2 snRNA transcript (*lsr1* gene). Distance in nucleotides to pairing partner in the predicted structure is plotted in red. Data for all replicates and mutants were merged. **(b)** Nucleotide composition of the indicated region within *lsr1*. **(c)** Secondary structure model of nascent U2 snRNA based on CoSTseq data from position 580 on. Data for wildtype and mutants were merged, n=20. **(d)** Schematic representation of unpaired and early pairing nucleotides during transcription (left) relative to the final folded state in the U2 snRNP (right).

In addition to RNA Pols I and II, we also capture co-transcriptional folding of RNA Pol III transcripts. Consistent with the other RNA Pols, overlaying CoSTseq data for the scR1 (signal recognition particle RNA) transcript on its secondary structure model^42^ indicates co-transcriptional formation of small stems (e.g. 9a, 9d, 10b), whereas long-range base pairs and more complex junctions (e.g. 5c, 6, 7, 8) are often in disagreement with nascent DMS reactivities (Fig. S2).

### Local base pairing of nascent rRNA differs radically from mature rRNA

Ribosomal RNA is highly structured with a well-established base pairing pattern in its mature state.^43^ CoSTseq enables us to evaluate local RNA folding patterns relative to RNA Pol positions *in vivo*, and to relate these changes to functional RNA processing. Additionally, our ability to visualize folding of the 5’ETS of pre-rRNA during transcription suggested we could learn more about the contribution of nascent RNA folding to the production of ribosomes. To ask how nascent rRNA folding potentially differs from mature rRNA, we complemented the CoSTseq dataset for nascent rRNA with DMS-MaPseq datasets specifically for mature rRNA (i.e. not polyadenylated). When comparing DMS signal across the whole rRNA locus by calculating PCC of DMS reactivities in windows across the transcript between biological replicates of the same sample type (DMS-MaPseq or CoSTseq), high reproducibility was evident (Fig. 3a). In contrast, we found extensive differences between nascent and mature rRNA, including the anticipated detection of structured nascent external and internal transcribed sequences (ETSs and ITSs) that are not present in mature rRNA. A few regions such as rRNA stem loops 21ES6cb and 63ES27a display high correlation suggesting they rapidly adopt their mature structure after exiting RNA Pol I (Fig. S3a). Overall, the extensive disagreement between CoSTseq and DMS-MaPseq indicates that nascent rRNA forms transient intermediate structures that undergo rearrangements during transcription *in vivo*.

**Fig. 3:**
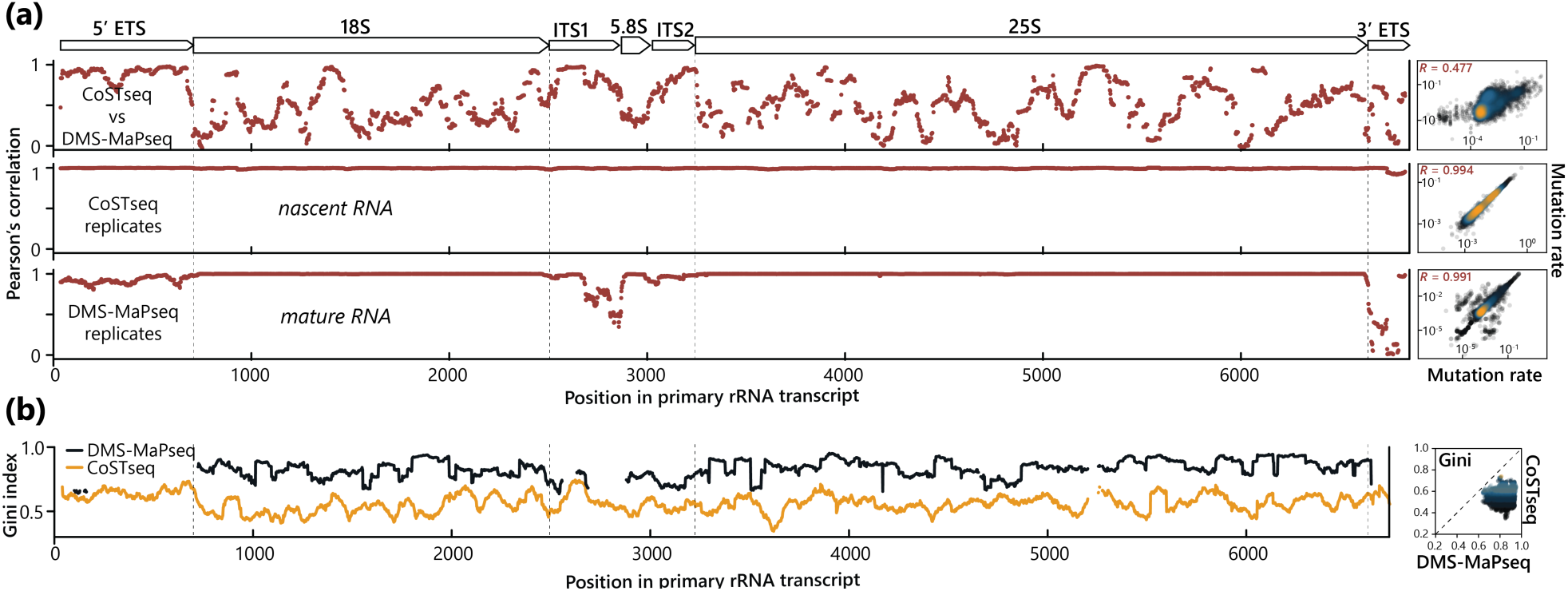
Co-transcriptional rRNA base pairing differs strongly from mature rRNA. **(a)** Pearson’s correlation coefficient at each nucleotide in sliding windows of size 50 across the primary rRNA transcript calculated between CoSTseq (n=3) and DMS-MaPseq (n=3) merged replicates (top), CoSTseq replicate one and CoSTseq replicate two (middle), DMS-MaPseq replicate one and DMS-MaPseq replicate two (bottom). The same correlation coefficients are replicated as scatter plots to the right and the bulk coefficient *R* is reported. **(b)** Gini index for CoSTseq or DMS-MaPseq data calculated for each nucleotide of the primary rRNA transcript in windows of size 80. The same Gini index values are replicated as scatter plots to the right. External Transcribed Spacer, ETS; Internal Transcribed Spacer, ITS.

If intermediate structures form and are remodeled later during ribosome biogenesis, we expect to find less defined structures when averaging RNA Pol I positions and this should decrease the Gini index. Indeed, we find a consistently lower Gini index in projected nascent rRNA signal, consistent with dynamic rearrangements and/or fluctuating structures at individual RNA Pol I positions (Fig. 3b). One possible explanation for this behavior would be that the first ≤400 nucleotides adjacent to RNA Pol I surveyed by CoSTseq lack their final base pairing partners. We analyzed this across the rRNA locus and found that only ∼14% of paired nucleotides are theoretically unable to engage in their final base pairing interactions within 150 nt after exit from RNA Pol I, because their partner will not have been added yet to the growing RNA chain. Thus, the local base pairs formed in nascent rRNA and detected by CoSTseq differ substantially from those attained in mature ribosomes despite the availability of final partner nucleotides. These data lead us to conclude that previously unknown, transient structural states of rRNA form on-pathway to ribosome maturation. Despite such rearrangements during transcription, projected DMS reactivities support previously-modeled secondary structures of the 5’-ETS^44,45^ (Fig. S3b). Importantly however, CoSTseq uniquely captures the 5’-ETS prior to U3 snoRNA binding and therefore provides new insight in the region of stems H4 and H5, which form distinctly more stable structures in our model than previously reported (Fig. S3c). This observation implies that some form of unwinding needs to take place before U3 can anneal. Since constrained structure prediction based on CoSTseq data recapitulated the known base pairs of the 5’ETS reasonably well, we performed structure prediction for the entire nascent rRNA transcript (Fig. S4a) and evaluated prediction accuracy across the locus (Fig. S4b). These predictions represent the most likely base pairs forming as nascent rRNA exits from RNA Pol I.

### Individual nucleotides display distinct kinetics of co-transcriptional base pairing

To understand the dynamics of co-transcriptional RNA folding in quantitative detail and specifically determine when base pairing interactions occur, we followed DMS reactivity as a function of transcription progression by combining 3’-end identity (to obtain RNA Pol position) and inferred methylation rates (to indicate the proportion of paired A/C nucleotides) for each nucleotide. Separate DMS reactivity profiles were calculated for most RNA Pol I positions by grouping reads according to 3’-end identity. Structural transitions were visualized by displaying profiles in the format of a DMS reactivity matrix (Fig. 4a). This analysis is supplemented by a projection of all overlapping DMS reactivity profiles (Fig. 4a, bottom); Nucleotides that display intermediate reactivities in projected DMS reactivity profiles often transition from high to low reactivities due to co-transcriptional base pairing.

**Fig. 4:**
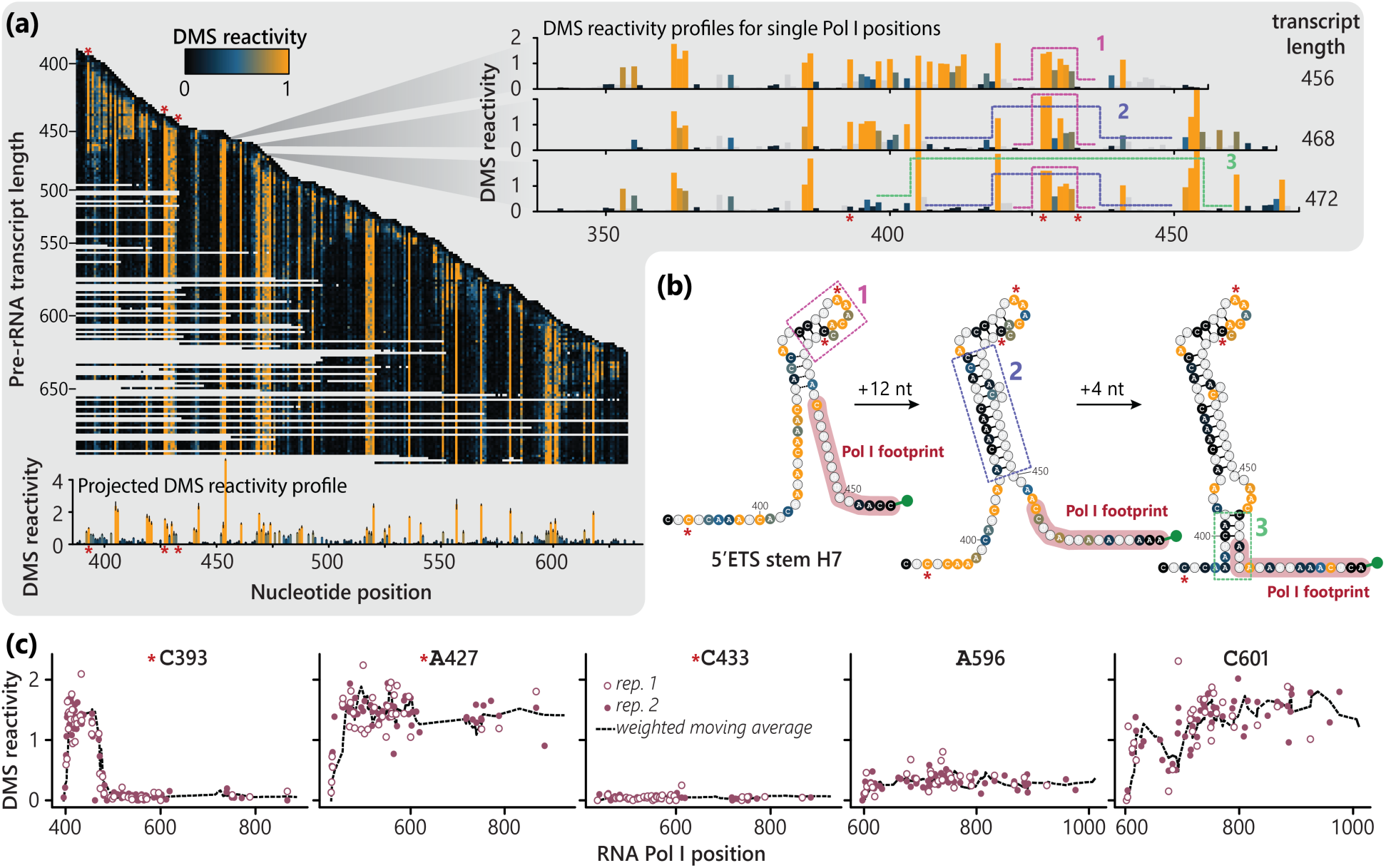
Nucleotide trajectories reveal distinct local rRNA base pairing behaviors, by associating RNA polymerase position with DMS reactivity for each nascent RNA read. **(a)** Co-transcriptional folding matrix for a section of the rDNA gene. Each row represents the DMS reactivity profile of the nascent pre-rRNA at the respective RNA Pol I position within the 5’-ETS, as indicated by the position of the profile along the *y*-axis. DMS reactivity profiles for three RNA Pol I positions (right) are enlarged and putative double stranded regions are marked by dashed lines. The projected DMS reactivity profile (bottom) represents the averaged DMS reactivities over all RNA Pol I positions, including U reactivities due to increased coverage. Error bars show standard deviation of at least three biological replicates at each position, n=5. Nucleotides highlighted with an asterisk correspond to the same nucleotides in (b) and (c). **(b)** Structure models of stepwise stem H7 formation in the 5’-ETS overlayed with DMS reactivity values from the enlarged profiles in (b). Nucleotides shaded in red are located within the RNA Pol I holoenzyme. Dashed rectangles show sequentially forming stem loop sections as transcription progresses, analogous to (b). **(c)** DMS reactivity trajectories for individual nucleotides; the first datapoint appearing on the x-axis is the position at which the nt was added to the growing nascent RNA chain. Dashed line represents the weighted moving average over 5 nt, signifying DMS reactivities as the nascent chain further elongates. Data for two representative replicates are shown.

To illustrate an example of co-transcriptional folding detected by CoSTseq, we followed the stepwise formation of stem H7 in the 5’-ETS (420-440 nt), showing that it forms in three steps starting with the segment close to the loop region immediately after exit from RNA Pol I (Fig. 4b). To study the timing of these rearrangements, we generated trajectories for each nucleotide that report DMS reactivity as a function of transcription progression (Fig. 4c). We found that individual nucleotides exhibit distinct modes of co-transcriptional behavior: delayed pairing (e.g. C393), no pairing (e.g. A427), immediate pairing (e.g. C433), or gradual pairing (e.g. C601).

Observation of the full co-transcriptional folding matrix for rRNA (Fig. 5a and Fig. S5a) suggest that there might be distinct types of nucleotide trajectories. These may signify base pairing characteristics which we might be able to quantify, and extract parameters describing the fundamental kinetics of co-transcriptional base pairing. Therefore, we first calculated a weighted moving average (as shown in Fig. 4c) for all trajectories with at least eleven data points and inferred reactivities for missing Pol positions through linear interpolation (Fig. 5a, left). Nucleotide trajectories were then classified through hierarchical clustering (see Methods) by applying a distance threshold and merging neighboring groups that appeared similar (Fig. 5a, right). This procedure yielded four major classes to which we assigned molecular identity based on their appearance (Fig. 5b): immediate pairing, delayed pairing, unpaired, and dynamic/mixed interactions, leaving only 1.8% of trajectories unclassified (Fig. 5c). We next returned to raw trajectories and sought to quantify base pairing of each nucleotide class by modeling DMS reactivity in response to transcription progression. We found that hyperbolic decay models describe DMS reactivity trajectories well (Fig. 5b) and allow us to extract fundamental parameters of co-transcriptional folding: when base pairing occurs (expressed as the distance at which base pairing is completed to 90%, *d*_0.9_), and the DMS reactivity saturation value (*r*_sat_), which informs on the final state of each nucleotide class. Unpaired nucleotides have a high *r*_sat_value and calculating *d*_0.9_ is not informative since it lies beyond the experimental window of observation (larger than ∼400 bp), consistent with remaining unpaired. The mixed class has an intermediate *r*_sat_value, which might indicate protein binding, tertiary structure formation, or interconversion with multiple structural states populated. Consistent with stable base pairing, the immediate pairing class has the lowest *r*_sat_value and these nucleotides reach *d*_0.9_within 25 bp of transcription, corresponding to an average pairing rate λ*_r_* of 48.5 kb^-1^. Delayed pairing (*d*_0.9_= 300 bp, λ*_r_*= 13.2 kb^-1^) occurs in a small number of cases (4.2%), likely when nucleotides are located at the left side of the base of large stem loops, as we previously showed for U2 snRNA (Fig. 2). Consistent with this model, delayed nucleotides appear in clusters across the pre-rRNA (Fig. S5b). In summary, quantitative description of DMS reactivity trajectories for pre-rRNA nucleotides revealed that ∼30% of nucleotides quickly engage in pairing interactions within 25 bp of transcription, which we consider rapid. In contrast, ∼45% of nucleotides remain unpaired during transcription; yet many of these may be in loops associated with stems, i.e. within structures (Fig 5b).

**Fig. 5:**
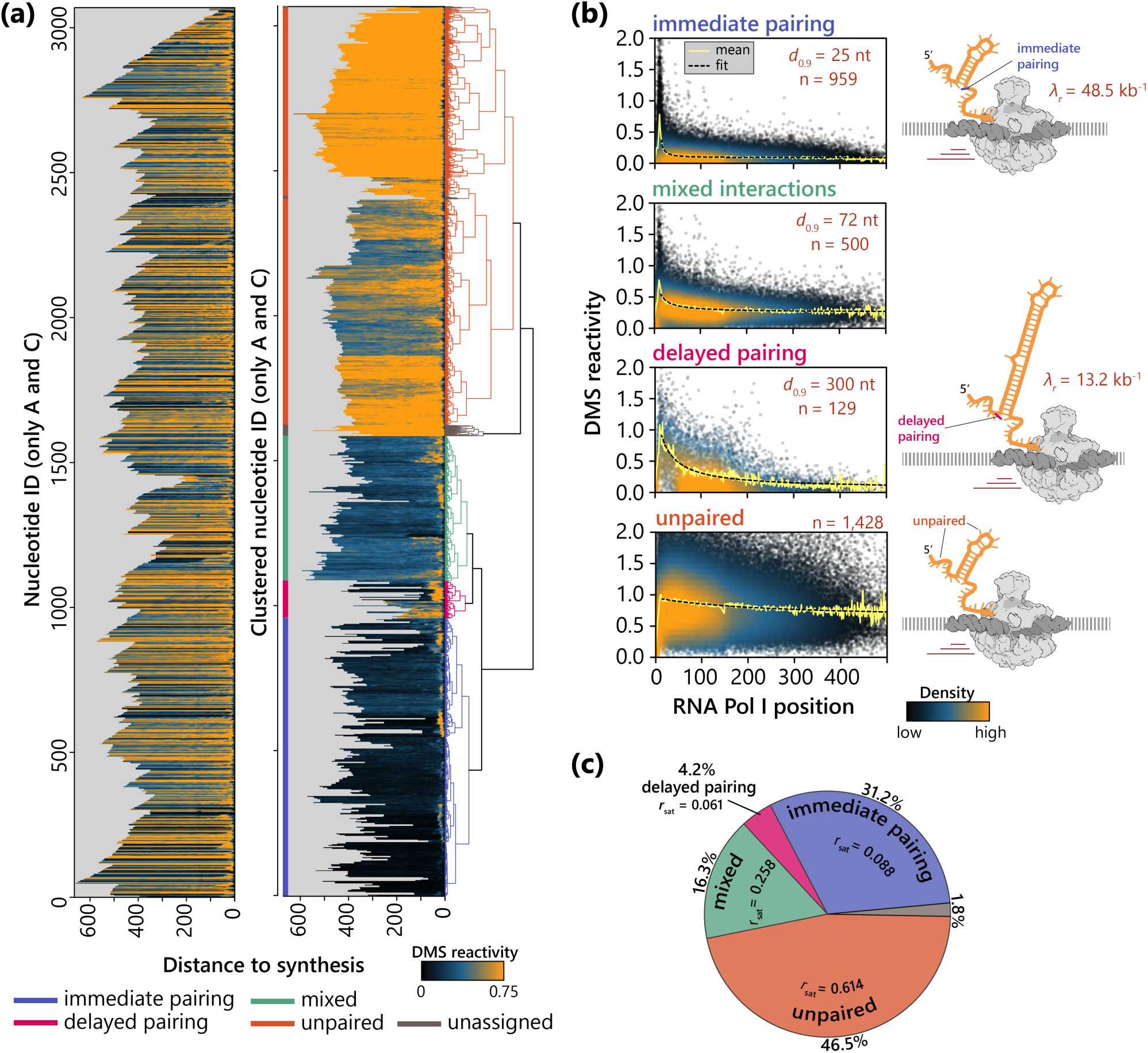
Classification and modeling of DMS reactivity trajectories reveals fundamental parameters of co-transcriptional base pairing. **(a)** Left: All interpolated DMS reactivity trajectories aligned at the 3’-end (Pol I position) and ordered according to nucleotide position. Right: Same trajectories as left but ordered according to position in the hierarchical clustering derived dendrogram. The dendrogram is colored according to class annotation. **(b)** Hyperbolic decay models describe all four classes of DMS reactivity trajectories. Trajectories for each class (raw data, not interpolated) were pooled and fitted. The model (black dashed line) describes the positional mean (yellow line) well. n: number of trajectories, *d*_0.9_: Pol I distance at which 90% of *r*_sat_ (reactivity at saturation; base paired state) are reached. **(c)** Relative abundance and *r*_sat_ of each nucleotide class.

### Helicase Dbp7 regulates rRNA base pairing during transcription

Almost half of pre-rRNA nucleotides remain unpaired during early transcription (Fig. 5), and we found that co-transcriptionally formed structures in pre-rRNA must later rearrange (Fig. 3). We therefore hypothesized that RNA helicases could play a role in maintaining nascent RNA flexibility already during transcription. In contrast to the transcription process, less is known about how nascent pre-mRNA structure is regulated by helicases, which act by a variety of mechanisms from limited, local perturbations of base pairing to the unwinding of base pairs.^26,27^ Indeed, ATP-dependent activity has been shown to alter rRNA and mRNA structure in yeast, possibly attributable to helicase action.^43^ Both Dbp3 and Dbp7 are RNA-dependent ATPases with sequence-independent helicase activity implicated in pre-60S rRNA processing.^46–50^ We also flagged the G4 quadruplex binding protein, Stm1, as potentially interesting and created fresh knock-out strains for these factors (*dbp3Δ*, *dbp7Δ* and *stm1Δ*). We generated CoSTseq and matching DMS-MaPseq datasets for all strains in three biological replicates and tested the effects of these deletions on large and small ribosomal subunit abundances. We find that Dbp7 has the strongest effect on the synthesis of the large subunit (Fig. 6a, Fig. S6a). Comparing DMS reactivity trajectories between wildtype and helicase-deficient strains, we observed marked differences in timing and extent of base pairing for several residues in *dbp7Δ* (Fig. 6b) but not *dbp3Δ* (Fig. S6b). We therefore plotted the DMS reactivity difference between wildtype and either *dbp7Δ*, *dbp3Δ* or *stm1Δ* at every nucleotide for every available RNA Pol I position (Fig. 6c). The overlay identifies and visualizes sites at which Dbp7 acts on nascent rRNA in a co-transcriptional manner. These include previously identified sites^49^ but, strikingly, also reveal that Dbp7 acts along the entire nascent transcript, while Dbp3 and Stm1 show no co-transcriptional RNA remodeling activity in our window of observation. To better quantify differences, we calculated the average reactivity difference between wildtype and mutants within each of the four major regions for the pre-rRNA transcript, using *spt4Δ* as a negative control (Fig. 6d). While *dbp3Δ* and *stm1Δ* show no changes in any of the regions compared to *spt4Δ* (Fig. S6d), *dbp7Δ* shows decreased base pairing particularly in the ITS region, where it was previously reported to facilitate a cleavage event.^49^ These findings enhance our understanding of how Dbp7 helicase activity is required across virtually the entire nascent pre-rRNA for ribosome biogenesis, maintaining the relative amounts of large and small subunits produced (see Fig. 6a and ref. 49).

**Fig. 6:**
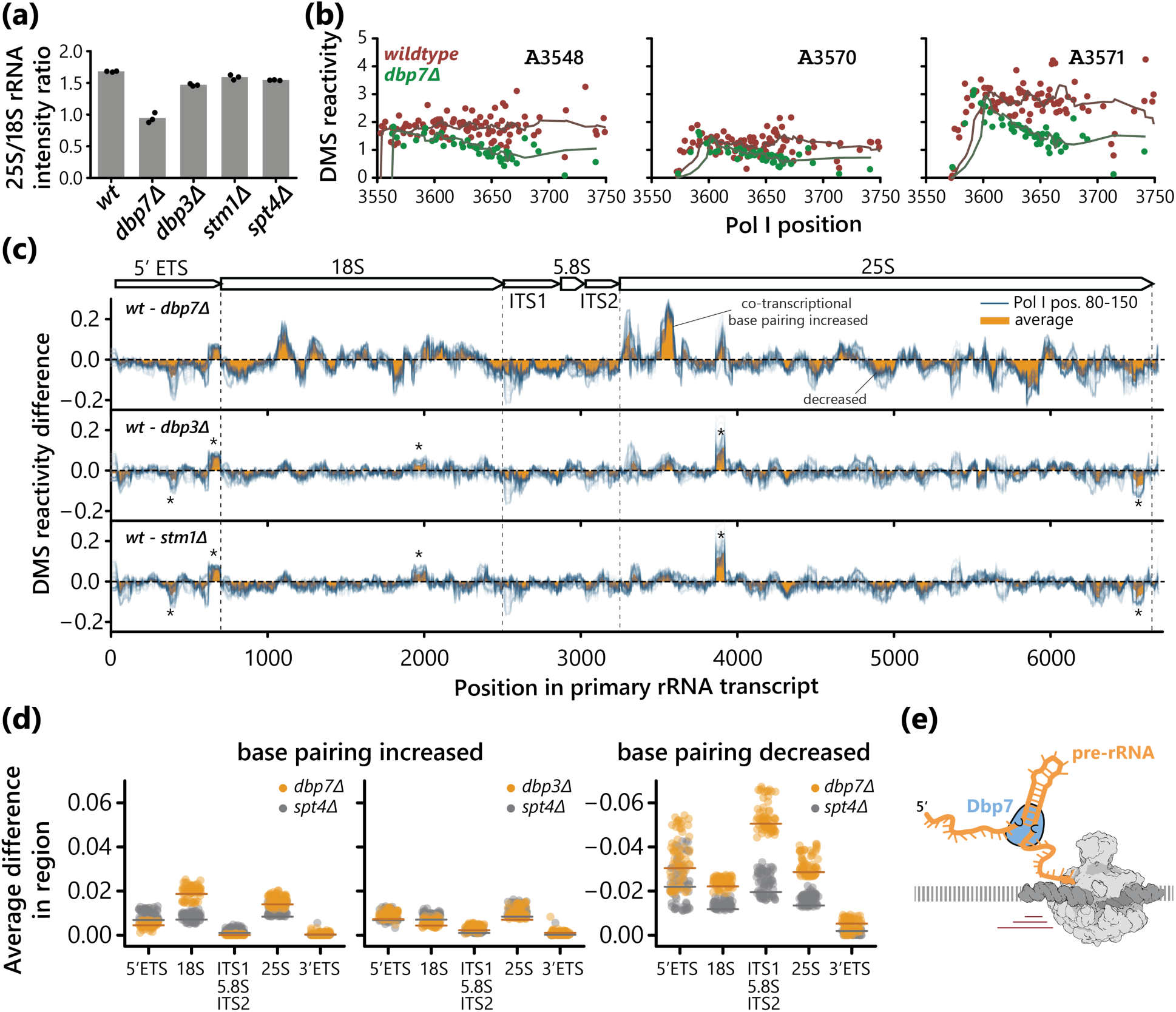
Helicase Dbp7 remodels local nascent rRNA structures as soon as they form. **(a)** Large to small ribosomal RNA ratio for wildtype and mutant strains assessed by agarose gel electrophoresis. **(b)** DMS reactivity trajectories for individual nucleotides. Solid line corresponds to the weighted moving average. Data for replicates were merged, wildtype: n=5, *dbp7Δ*: n=3. **(c)** DMS reactivity difference between wildtype and helicase mutants. Each blue line represents data from reads where RNA Pol I was between 80 and 150 nucleotides past the nucleotide of interest, one line for each relative RNA Pol I position. The average difference between all RNA Pol I positions 80-150 is shown in orange. Asterisks indicate regions that appear different in all conditions and can therefore be considered as background. Data for replicates were merged, wildtype: n=5, *dbp7Δ*: n=3, *dbp3Δ*: n=3. **(d)** Regional average of average (across Pol I positions 80-150) DMS reactivity difference between wildtype and each indicated mutant, using *spt4Δ* as a control. **(e)** Schematic showing the co-transcriptional association of Dbp7 helicase with nascent pre-rRNA as it emerges from RNA Pol I.

### Local nascent RNA base pairing determines mRNA structure

Base pairing is a major contributor to mRNA stability, and secondary structure in the cytoplasm correlates with protein expression.^14,51^ Yeast pre-mRNA undergoes co-transcriptional splicing and mRNP formation. To what extent mRNA structure is remodeled prior to export and translation – or affected by translation – remains unclear. To test the hypothesis that pre-mRNA base pairing is remodeled after transcription, we compared nascent and mature RNA structure probing data (Fig. 7a). We observe remarkable agreement when comparing nascent with its mature states for almost all transcripts (median PCC of 0.90 per transcript, Fig. 7b). In addition, Gini indices for individual transcripts are also highly correlated and differ slightly but significantly only for intron-containing transcripts (Fig. 7c). This agreement suggests that no significant remodeling takes place post-transcriptionally and that the base pairs forming after the RNA emerges from RNA Pol II are final. Supporting this conclusion, unconstrained structure predictions with smaller maximum distance between base pairs describes our mRNA DMS-MaPseq data better, as shown by calculating the area under the receiver operator characteristic (AUROC) curve for a range of maximum base pair distances for all well-covered mRNA transcripts (Fig. S7a). Analysis of previously published icSHAPE data from fractionated human cells^52^ similarly shows strong structural agreement between chromatin-associated and cytoplasmic RNA (Fig. S7b) suggesting that concordance of nascent and mature conformations is a conserved property across organisms. To further test in our system whether base pairing might depend on helicases or RNA Pol II elongation, as suggested previously^22^, we compared CoSTseq and DMS-MaPseq signal between wildtype and the elongation-defective *spt4Δ* strain. Our previous analysis had shown no effect on nascent RNA reactivity within RNA Pols I and II (see Fig. 1e-f), but changes in overall nascent or mature mRNA folding are not ruled out by that analysis. However, such changes were not detectable by CoSTseq (Fig. S7c), indicating that altered elongation properties through lack of Spt4 might not affect base pairing during RNA Pol II transcription in our system.

**Fig. 7:**
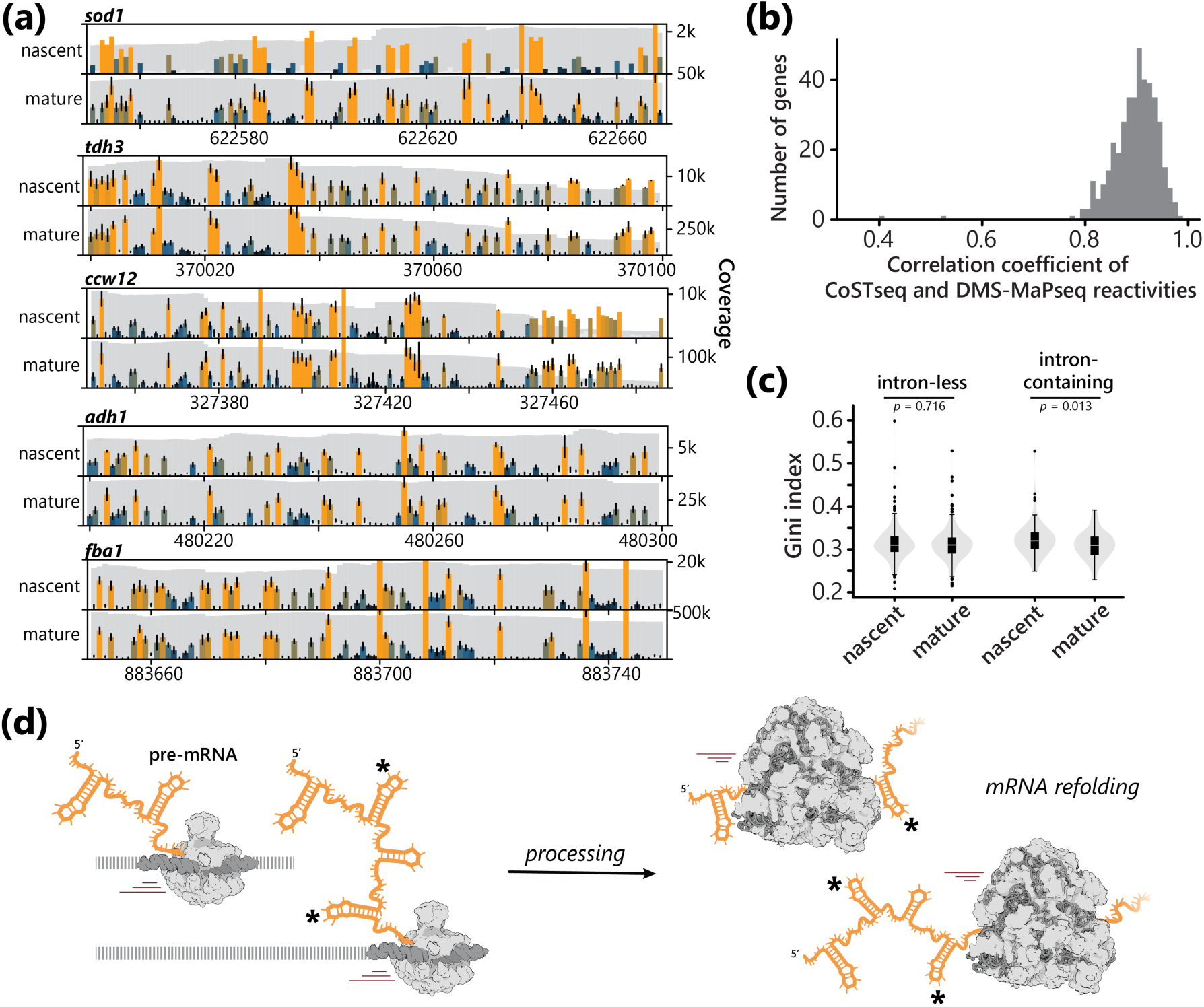
Nascent pre-mRNA and mature mRNA structures are mostly indistinguishable. **(a)** DMS reactivity profiles from CoSTseq (nascent RNA) and DMS-MaPseq (mature RNA) reads. Light gray bars indicate G/U nucleotides. Bars at A/C nucleotides are colored according to normalized signal. Coverage is indicated in dark gray. Error bars signify standard deviation across at least three biological replicates. Bars without error bars do not have enough coverage in multiple replicates to calculate a biological error. Data from wildtype and all mutants (*spt4Δ*, *dbp7Δ*, *dbp3Δ*, *stm1Δ*) with three replicates each were included to calculate profiles, CoSTseq: n=15, DMS-MaPseq: n=15. **(b)** Distribution of Pearson’s correlation coefficients of DMS reactivities between nascent and mature RNA calculated for individual transcripts. Data for all replicates and mutants were merged as in (a). **(c)** Gini index for individual transcripts separated by transcript class. Data for all replicates and mutants were merged as in (a). P-values for comparing nascent with mature RNA Gini indices are calculated according to two-sided Mann-Whitney U test. **(d)** Schematic representation of co-transcriptional pre-mRNA folding and co-translational mRNA refolding, which are similar based on the comparison between CoSTseq and DMS-MaPseq data.

To test more rigorously whether any of the deletion strains display changes in mRNA base pairing, we sought to compare wildtype DMS-MaPseq to mutant data, but tools for statistically robust and well powered comparisons of mutational structure probing data have so far only been developed for gene-specific structure probing datasets. To analyze all sequenced transcripts, we developed a novel statistical approach based on a previously reported Bayesian hierarchical model^53^, which allowed us to perform pairwise comparisons for genome-wide mutational probing datasets (HDProbe, Fig. S7d). This framework requires minimal user input as all parameters that influence inference are learned from the data (see Methods). Using HDProbe, we identified significantly differentially modified nucleotides in DMS-MaPseq data, comparing the wildtype to each of the mutants. Surprisingly, *dbp7Δ* (but not the other mutants) displays many significant changes in polyadenylated mRNA base pairing (Fig. S7e). Nucleotides with significant differences (p < 0.05) tend to co-occur in the same transcripts particularly towards the 5’-end (Fig. S7f), indicating that the helicase may act on mRNA in addition to its known role in rRNA processing, potentially prior to export.

## Discussion

RNA and protein share the property of attaining specific folded states, which can enable critical cellular functions that can be catalytic, regulatory, and/or related to the formation of higher order complexes. Co-translational folding of nascent proteins is a familiar concept^54^, but analogous mechanisms are unknown for RNA. Here, we address the origin of folded RNA structures in eukaryotes by identifying the first steps of base pairing that take place co-transcriptionally *in vivo*. Our data reveal that the nascent RNA’s path through eukaryotic RNA polymerases is unfavorable for intramolecular base pairing, which begins as the nascent RNA emerges from the RNA polymerase exit channel. Co-transcriptional pre-mRNA structure appears to be generally indistinguishable from mRNA structure, indicating that local folding is dominant over possible longer-range interactions. In contrast, nascent pre-rRNA folds into transient structures across the rDNA locus that differ strongly from mature rRNA. We find that the helicase Dbp7 has unexpectedly broad effects on local base pairing along the full length of nascent pre-rRNA, by maintaining open states for some nucleotides as they exit Pol I and favoring folded states elsewhere. This role of Dbp7 appears to be remarkably similar to the action of protein chaperones in preventing premature folding of nascent peptides until later motifs are translated and integrated into the correct folding pattern.

These findings were made possible by the establishment of CoSTseq, a method that rigorously identifies the catalytic sites of elongating RNA polymerases and – upon DMS probing – yields reproducible, quantitative nascent RNA base pairing information for each step of transcription along endogenous eukaryotic genes. Remarkably, the obtained datasets support tracking of individual nucleotides for up to 400 elongation steps. These nucleotide trajectories can be classified into four main base pairing behaviors (see Fig. 5) that support a model whereby nascent RNA undergoes extremely rapid structure formation. Immediate pairing may start as soon as a nucleotide exits from RNA polymerase (*d* = 16 bp, see Fig. 1e-f, Fig. S1d-e) and finish on average after *d*_0.9_ = 25 bp from synthesis. Persistently unpaired categories are expected for loop regions of quickly formed, stable stem loops, or for flexible regions. Nucleotides belonging to the delayed category (*d*_0.9_ = 300 bp) likely signify a delay until the partner nucleotide is synthesized or perhaps by another regulatory step imposed by RNA modification or binding by protein. Using transcribed distance (bp) as a proxy for time, we calculated rates of folding for the rapid and delayed groups of RNA nucleotides as 48.5 and 13.2 kb^-1^, respectively. These metrics should be helpful for future computational approaches that seek to derive RNA folding pathways and potential interactions with other co-transcriptional events, such as RNA processing and modification.

Our data illuminate the initial step of ribosome biogenesis, identifying previously unknown, transient structural states that are markedly distinct from mature rRNA. These dynamic states are detected even before U3 snoRNA annealing can take place. Our *in vivo* results complement findings from cell-free systems that show dynamic association of chaperones guiding nascent RNA folding right after its synthesis.^55–57^ Through detection of dynamically changing base pairing status at the level of single RNA nucleotides and as a function RNA Pol I position, we also corroborated *in vitro* data showing that nascent RNA can be metastable.^58^ In addition, our study shows *in vivo* that the RNA helicase Dbp7 remodels nascent rRNA as it exits RNA Pol I, shedding new light on previous studies that established several roles in pre-60S processing. Interestingly, Dbp7 regulates base pairing between pre-60S and snR190 snoRNA near the eventual peptidyl transferase center (PTC); this observation highlights the importance of Dbp7 in modulating RNA-RNA interactions in cis and trans.^50^ Although Dbp7 helicase activity is not required for 2’-O-methylation or pseudouridylation, differences between wildtype and *dbp7Δ* may result in changes of nascent rRNA base pairing with snoRNAs that guide other modifications.^49^ Dbp7 plays a role in the recruitment of the early pre-60S maturation factor uL3, with which it appears to make a relatively stable complex on the 3’ end of the 25S pre-rRNA.^49^ Our CoST-seq data reveal an overall decrease of structure formation across this region of the 25S sequence in the absence of Dbp7, consistent with the model that Dbp7, uL3 and other factors act to compact this region prior to PTC maturation. Importantly, the previous study also discovered an independent function of Dbp7 in facilitating exonucleolytic processing at the A2, A3 and B1L/S sites within ITS1.^49^ Cleavage at the A3 site by MRP RNase is by weak sequence specific recognition of single-stranded RNA adjacent to several conserved stem loops,^59^ which we observe to be formed co-transcriptionally according to our CoSTseq data (see Fig. S4). The decrease in DMS reactivity across ITS1, 5.8S and ITS2 in the *dbp7Δ* strain indicates that co-transcriptional remodeling by Dbp7 may make this set of pre-rRNA cleavage sites available to ribosomal RNA processing factors and endonucleases.^60^ Our evidence for widespread remodeling of rRNA by Dbp7 may explain why 18S rRNA was also reduced in the absence of Dbp7 activity^49^ and raises the question of how Dbp7 is recruited to the rDNA locus. In conclusion, our findings demonstrate the functional roles helicases can play in gene expression by remodeling co-transcriptional RNA folding in coordination with RNA processing.

For mRNA, remodeling of initially formed base pairs appears to be the exception. Because translation elongation occurs on similar timescales as transcription^61^, it is conceivable that local mRNA base pairs re-fold after exit from ribosomes in the same way in which they initially arose during transcription. This notion is consistent with experiments in zebrafish that showed different structures in coding regions but not UTRs when comparing *in vivo* with *in vitro* re-folded mRNA structure, suggesting that maintenance of local base pairing in the cytoplasm is dependent on translation.^62^ Taken together, our data support an RNA-centric model of RNA maturation and structure acquisition, whereby intramolecular base pairing upon exit from RNA polymerases determines the final structural state. Protein chaperones, such as helicases, are abundant and varied in their functions, with many mysteries to be solved regarding mechanism.^26,27^ As we have seen, some helicase activities have been associated with specific RNA processing steps, as had Dbp7. Yet, CoSTseq reveals effects of Dbp7 on pre-mRNA. Thus, it seems that helicases may guide the folding of multiple RNA classes. More information is needed to understand how helicases are targeted to their substrates.

These findings change how we imagine nascent RNA, the substrate of many processing steps. Nascent RNA is often represented as a string-like, linear molecule with sequence accessible for recognition by trans-acting factors that recruit diverse machineries. Many computational approaches to finding enhancing or repressive sequences in nascent RNA assume availability to the ∼1,000 RNA binding proteins expressed by eukaryotes. On the contrary, the demonstration by CoSTseq of immediate nascent RNA folding leads to the expectation that nascent RNA is structured and that cis elements may often be sequestered in competing structures. This concept is consistent with observations that RNA folding during transcription is an important part of its function, as it can regulate co-transcriptional processing events.^8^ So far, single-nucleotide and RNA Pol resolved data were limited to bacteria with different RNA processing pathways or limited to *in vitro* studies.^16,18,63^ As recently shown in prokaryotes^17^, it appears that also eukaryotic RNA folding and on-pathway structural transitions occur while RNA is still attached to chromatin. Our example of the co-transcriptional pairing of U2 snRNA provides a counterexample, whereby a relatively long sequence (440 nt) can refrain from base-pairing and then become immediately captured into a large stem loop when the complementary sequence is synthesized. We speculate that RNA folding patterns in eukaryotes may depend on the large constellation of RNA binding proteins^64^ that are known to regulate highly complex co-transcriptional RNA processing reactions. If so, tissue identity and/or signaling in response to environmental cues could act through widespread roles of nascent RNA folding as an effector mechanism of gene regulation.

### Limitations of the study

Like all RNA chemical probing methods where modification rate is inferred from mutations or RT fall-off, detection of specific transcripts using CoSTseq depends on the number of reads available. Since the majority (∼60%) of CoSTseq reads align to rDNA, the remaining reads are spread across all other expressed genes, making sequencing depth the limiting factor specifically for the analysis of Pol II and Pol III transcripts. In addition, the interpretation of folding kinetics must be conditioned on the experimental requirements, i.e., allowing time for transcriptional run-on and DMS modification reactions. It is possible that RNA structural re-arrangements occur during this time frame while Pols are already stopped. Offsetting this potential caveat is the stochasticity with which biotin incorporation stops elongation complexes (e.g., at the beginning versus at the end of the 2 min run-on). If there are rearrangements (occurring on yet unknown timescales), the DMS signal corresponds to a population average across several thousand observations (reads) in our dataset for each bp of transcription. In the future, this could be addressed by employing structural ensemble deconvolution tools^10–13^ for reads that align to single Pol positions; currently their high read depth requirement is limiting. Finally, detection of base pairing depends on the availability of each nucleotide to DMS methylation. By comparing *in vivo* with *in vitro* DMS probing, many prior studies have reported general concordance of DMS profiles if the RNA exhibits the same folding pattern. However, due to the presence of cellular constituents, especially intermediate DMS reactivities cannot always be unambiguously assigned to a base pairing state.

## Methods

### Biotin-NTP run-on, DMS treatment and RNA extraction

Cultures from individual colonies of *S. cerevisiae* were grown in liquid YPAD medium to an OD_600_ around 0.6 at 30 °C with 200 rpm shaking. 2.8 ml culture were removed, spun down, and washed with cold PBS. The pellet was resuspended in 10 ml cold 0.5% sarkosyl and incubated on ice for 20 min. Cells were then pelleted at 400×g. Run-on reactions were performed in a 300 µl volume with 25 µM biotin-11-CTP (Jena Biosciences), 125 µM each ATP, UTP and GTP, 0.5% sarkosyl, 20 mM Tris⋅HCl pH 7.4, 200 mM KCl, 5 mM MgCl_2_ and 2 mM dithiothreitol at 30 °C for 2 min with mild shaking. 200 µl prewarmed 2.5x structure probing buffer (750 mM Bicine⋅KOH pH 8.0, 500 mM CH_3_CO_2_K pH 8.0, 12.5 mM MgCl_2_) and 25 µl DMS were added and incubated for 4 min at 30 °C with mild shaking. The reaction was stopped by the addition of 1 ml ice cold stop buffer (20 mM Tris⋅HCl pH 7.4, 200 mM KCl, 4.29 M 2-mercaptoethanol, 50% isoamyl alcohol). Cells were pelleted at 3,500×g and washed with stop buffer without isoamyl alcohol, then resuspended in 600 µl lysis buffer (45 mM NaOAc, 6 mM EDTA, pH 5.5) with 40 µl 20% SDS added after resuspension. The mixture was then incubated at 65 °C for 30 s with vigorous shaking, added to 650 µl hot (65 °C) acid-phenol:chloroform:IAA 125:24:1, pH 4.5 and incubated for 2 min at 65 °C with shaking. Samples were placed on ice for 5 min, then spun at 20,000×g at room temperature. The aqueous layer was subsequently again extracted with acid-phenol, then with chloroform and RNA was precipitated with isopropanol.

### CoSTseq library preparation

Approximately 80 µg of input RNA were incubated with Dynabeads MyOne Streptavidin C1 (Invitrogen) according to the manufacturer’s recommendations. The supernatant of the pulldown was retained for mRNA isolation. Beads were washed twice with high salt buffer (2 M NaCl, 50 mM Tris⋅HCl pH 7.4, 0.5% Triton X-100), once with binding buffer (300 mM NaCl, 10 mM Tris⋅HCl pH 7.4, 0.1% Triton X-100), and once with low salt buffer (5 mM Tris⋅HCl pH 7.4, 0.1% Triton X-100); 500 µl each. Nascent RNA was extracted from beads twice using Trizol and chloroform and resuspended in 3.75 µl H_2_O. Reverse transcription was performed in a 10 µl volume with an optimized KCl based buffer^65^ made of 75 mM KCl, 3 mM MgCl_2_, 50 mM Tris⋅HCl pH 8.3 and 0.5 µl RNase inhibitor using the GsI2c-MBP fusion protein (commercially TGIRT-III), which was purified as described previously^29^. RT reactions were pre-incubated with 100 nM annealed DNA/RNA heteroduplex adapter with a single G overhang to capture the biotin-C at the 3’-end (GTGACTGGAGTTCAGACGTGTGCTCTTCCGATCTTG, `rArArGrArUrCrGrGrArArGrArGrCrArCrArCrGrUrCrUrGrArArCrUrCrCrArGrUrCrArC/3SpC3/`) for 30 min at room temperature. Then 1.25 µM dNTP were added and the reaction was incubated at 25 °C for 10 min, 42 °C for 10 min, 50 °C for 10 min, 55 °C for 10 min, 60 °C for 30 min, 65 °C for 20 min, and 75 °C for 15 min. 1 µl of RNase H (NEB) was added and the sample incubated at 37 °C for 30 min, following clean-up using MinElute PCR Purification kit (Qiagen). The 5’-end adapter (`/5Phos/NNNNNNNAGATCGGAAGAGCGTCGTGTAGGGAAAGAGTGT`/`3SpC3/`) was first adenylated using Mth RNA Ligase (NEB), purified using Oligo Clean & Concentrator (Zymo Research) and then ligated to cDNA using Thermostable 5’ App DNA/RNA Ligase (NEB) according to the manufacturer’s instructions. Adapter ligated cDNA was again cleaned using spin column purification. PCR was carried out with KAPA HiFi HotStart polymerase (Roche) using Illumina TruSeq index primers (i5: `AATGATACGGCGACCACCGAGATCTACACXXXXXXXXACACTCTTTCCCTACACGACG`, i7: `CAAGCAGAAGACGGCATACGAGATXXXXXXXXGTGACTGGAGTTCAGACGTGTG`) for 16-20 cycles. Final libraries were size selected in the 200-1,000 bp range by cutting from a polyacrylamide gel and sequenced on Illumina NovaSeq machines in 2×150 paired-end mode.

### DMS-MaPseq library preparation

RNA was precipitated from the supernatant of the biotin pulldown using ethanol. 50 µg were used as input for the Dynabeads Oligo(dT)_25_ mRNA purification kit (Invitrogen) and mRNA was purified according to the manufacturer’s instructions. RNA was then fragmented using 10 mM ZnCl_2_ at 94 °C for 55 s and precipitated using isopropanol. End repair using T4 PNK was performed according to the manufacturer’s instructions. Fragmented mRNA was then size selected by differential silica binding with RNA Clean & Concentrator-5 spin columns (Zymo Research) using 33% ethanol. RT was carried out as described above, except the DNA strand contained a random overhang (N) instead of a G. Adapter ligation, PCR and sequencing were performed as described above. Mature rRNA DMS-MaPseq was carried out by fragmentation and library preparation without any prior enrichment steps.

### Knockout strain construction

*S. cerevisiae* BY4741 (ATCC 4040002) knockout strains were constructed by kanamycin cassette replacement. Amplicons with homology around start and stop codon of the target gene were generated from pFA6a-KanMX6 by PCR and transformed into *S. cerevisiae.*^66^ Resulting kanamycin resistant colonies were screened by PCR and confirmed by Sanger sequencing for correct gene replacement. Loss of target gene expression was confirmed in DMS-MaPseq and CoSTseq datasets.

### CoSTseq data processing

The modular CoSTseq data analysis pipeline, implemented including all custom sub modules in Snakemake, is available at https://github.com/NeugebauerLab/CoSTseq. Raw paired-end reads were quality filtered and trimmed with fastp^67^ and then aligned to the reference genome with STAR^68^, without generating new splice junctions. Ribosomal RNA was separated based on alignment coordinates and both mRNA and rRNA were deduplicated separately with UMICollapse^69^. Mutation counts and coverage for each nucleotide were extracted using a custom Python implementation that considers the following read filtering steps: Allowing maximum one clipped base at the 3’-end (and registering as a mutation if clipped); allowing maximum five clipped bases at the 5’-end; allowing maximum 15% of the nucleotides in a read to be mutated; a nucleotide must have a quality score of at least 20 to be considered a mutation; deletions are mapped as mutations as previously described^70^. For regular DMS-MaPseq (mRNA), unlimited clipping at the 5’-end was allowed if the clipped bases were at least 80% A; five clipped bases were allowed at the 3’-end. Mutation rates were calculated as the ratio of mutations to coverage; empirical pseudo-counts were added to avoid overinterpretation of data from low coverage sites using a simple beta distribution model:

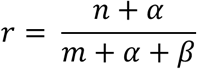

where *n* is the number of mutations and *m* is the coverage. Beta distribution parameters α and β were estimated from the distribution of all site-specific unadjusted mutation rates using the method of moments. Normalized DMS reactivity values were obtained as described previously^71^, by dividing by the mean of the top 10% of mutation rates after values above a threshold *max*(1.5 ∗ *IQR*, *P*_0.9_), where *P*_0.9_ denotes the 90^th^ percentile, were removed.

### Genome-wide pairwise comparison of mutation rates using HDProbe

HDProbe is an R package that compares nucleotide-specific mutation rates in chemical probing RNA-seq datasets. It adapts the statistical modeling strategy of a separate R package (bakR) to increase its statistical power.^53^ In short, HDProbe starts by filtering out low coverage sites to reduce the multiple-testing burden. Then, it infers the linear relationship between the logarithm of the replicate-to-replicate variability in the logit mutation rate estimates for a particular site and the logarithm of the coverage of that site. This linear relationship is used as a prior to regularize site-specific replicate variability estimates. A z-score is then calculated as the ratio of the difference in logit mutation rate estimate and the total regularized replicate variability. This z-score is then compared to a standard normal distribution to calculate a p-value, which is subsequently multiple-test adjusted using the Benjamini-Hochberg method.^72^

### Structure prediction

For structure prediction based on experimental DMS reactivity values, we used the rsample function of the RNAstructure package^73^ with the options -t 303.15 and --DMS. Normalized DMS reactivities larger than one were set to one. Pairing probabilities and the most likely pairs were calculated using the ProbabilityPlot and MaxExpect functions, respectively. For visualization, nucleotide coordinates were calculated from dotbracket files using the RNApuzzler algorithm.^74^ For AUROC computations with DMS-MaPseq data, the minimum free energy structure was predicted using the Fold function from RNAstructure and the option -md was varied.

### Nucleotide trajectory classification and modeling

Co-transcriptional folding matrices are composed of stacked DMS reactivity profiles for an RNA of increasing length, such that each row represents a DMS profile at a specific polymerase position. To generate these matrices from genome-wide CoSTseq data, a mutational and a coverage matrix were initialized. We then iterated through aligned read pairs and determined the 3’-end (Pol) position from the alignment coordinate of the Illumina R2 end. Reads with more than one clipped base at the 3’-end were discarded. Alignment coordinates of reads with one clipped base were shifted accordingly. The same read quality and mutational filters as described above were applied, and if the read passed, mutational and coverage matrices were updated based on mutation and alignment position of each read, respectively. To obtain DMS reactivity matrices, the ratio of mutation and coverage matrices was normalized as described above.

Nucleotide trajectories correspond to columns in the reactivity matrix. Due to sequencing depth limitations and run-on conditions with only biotin-CTP, matrices are sparsely populated. We therefore interpolated missing values between the initial addition of the nucleotide to the chain (zero coordinate) and the maximum observed Pol position using a weighted moving average, where the weight of each value decreases with its distance to the nearest point. This allows for preferential emphasis on regions with many data points, making the moving average less sensitive to outliers. To classify interpolated DMS reactivity trajectories, a hierarchical clustering model was used. Uniformity in length for hierarchical clustering was ensured by padding trajectories to the maximum observed trajectory length with a value of zero. Clustering was performed on the padded trajectories using the Ward linkage method with a Euclidean distance metric.^75^ The resulting dendrogram was trimmed at varying thresholds to create flat clusters, and an optimal threshold of 40 was determined empirically by assessing the resulting clusters for uniformity and distinctiveness. This yielded several neighboring classes which we merged based on appearance. Classes that were not neighbors were never merged. Classes with less than ten trajectories were labeled as unassigned.

Based on class assignments, original trajectories (without interpolation) were aligned at their zero coordinate, generating data-rich meta trajectories for each class. We then fitted a hyperbolic decay model describing the DMS reactivity *r* as a function of Pol position *d*

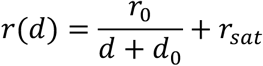

where *r*_0_ and *d*_0_ control the magnitude and onset of decay, and *r_sat_* represents the DMS reactivity at saturation. Fits were performed using the least squares method and evaluated by comparing to the positional mean of the meta trajectories. The DMS reactivity at which a fraction *f* (set to 90%) of the base pairing reaction is complete was calculated according to

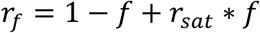

giving the corresponding Pol position with:

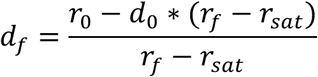

The Pol position-dependent rate of decrease in DMS reactivity λ*_r_*(*d*) is given with:

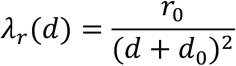

A nucleotide with DMS reactivity of zero is base paired and a nucleotide with DMS reactivity of one is unpaired, thus the rate of decrease in DMS reactivity can be considered equivalent to a base pairing rate. We report the average rate λ*_r_* between *d*_-_ = 15 (exit from Pol) and *d*_2_, = *d*_f=0.5_ (50% paired) according to:

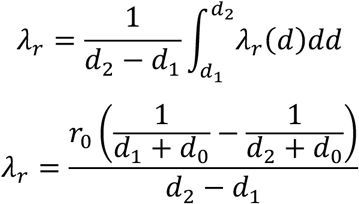

## Data availability

CoSTseq and DMS-MaPseq raw data are available through the National Center for Biotechnology Information (NCBI) under GEO accession number GSE254264. Prior to publication, GEO data can be accessed using a token which is available upon request.

## Code availability

Code for CoSTseq and DMS-MaPseq data processing, as well as downstream analysis is publicly available at https://github.com/NeugebauerLab/CoSTseq. HDProbe is publicly available at https://github.com/isaacvock/HDProbe.

## Acknowledgements

The authors would like to thank Sarah Woodson, Tucker Carrocci, Jackson Gordon and Paulina Podszywałow-Bartnicka for helpful discussions, and Pernille Bech for assistance with library preparation. This work was supported by the National Institutes of Health R01GM112766 (to K.M.N), NIH R01GM137117 (to M.D.S.), T32GM67543 (I.W.V.), and a predoctoral fellowship from the American Heart Association (908949 to L.S.). Data acquisition at Yale Center for Genomic Analysis was supported by the National Institute of General Medical Sciences of the National Institutes of Health under Award Number 1S10OD030363-01A1. This work is solely the responsibility of the authors and does not necessarily represent the official views of the NIH.

