## Supplementary Information for "Rapid folding of nascent RNA regulates eukaryotic RNA biogenesis"

**Title:**

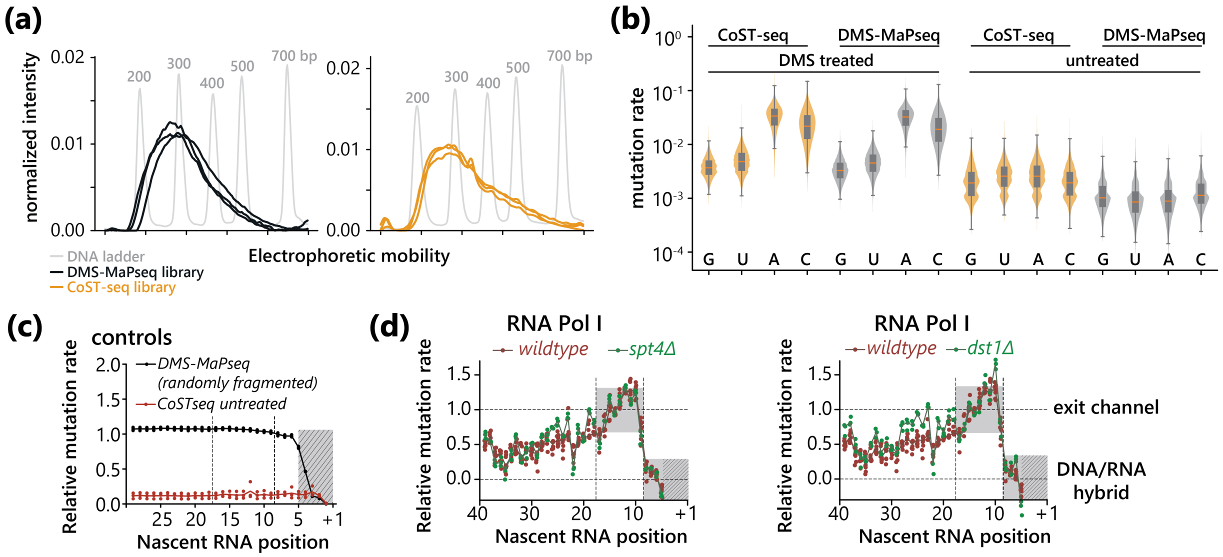


**Fig. S1** *(Related to Fig 1)***:** CoSTseq detects base pairing of nascent RNA with similar signal-to-noise ratio to DMS-MaPseq. **(a)** Capillary electrophoresis showing size distributions of DNA libraries for CoSTseq and DMS-MaPseq. **(b)** Mutation rate distributions for individual nucleotides in CoSTseq and DMS-MaPseq experiments, with or without DMS addition. Data from wildtype and mutants (*spt4Δ*, *dbp7Δ*, *dbp3Δ*, *stm1Δ*) with three replicates each were included to calculate mutation rates, CoSTseq DMS treated: n=15, DMS-MaPseq DMS treated: n=15, CoSTseq untreated: n=3, DMS-MaPseq untreated: n=2 **(c)** Mutation rates for nucleotides relative to the RNA 3'-end (+1) calculated from wildtype merged DMS-MaPseq data sets (n=3) and CoSTseq untreated data sets (n=2). DMS-MaPseq data is normalized to the mean of positions 50 to 140 where mutation rates are constant; CoSTseq untreated data is normalized in the same way with respect to CoSTseq DMS treated samples. **(d)** Mutation rates for nucleotides close to the RNA 3'-end (Pol I active site, +1) calculated from all reads aligned to rDNA. Mutation rate signal is normalized between the mean of positions 6-8 (set to zero) and 12-18 (set to one). Dots indicate individual biological replicates; solid line represents the median. Wildtype: n=8, *spt4Δ*: n=3, *dst1Δ*: n=3.

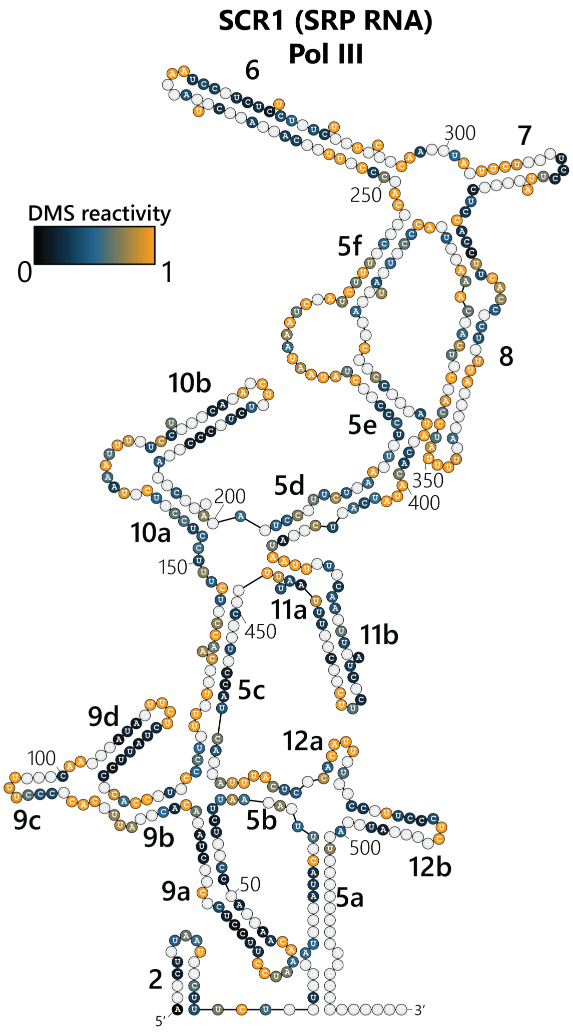


**Fig. S2** *(Related to Fig 2)***:** Secondary structure model of yeast SCR1 from Van Nues & Brown[48] overlaid with nascent RNA DMS reactivity. CoSTseq data for wildtype and mutants merged, n=20.

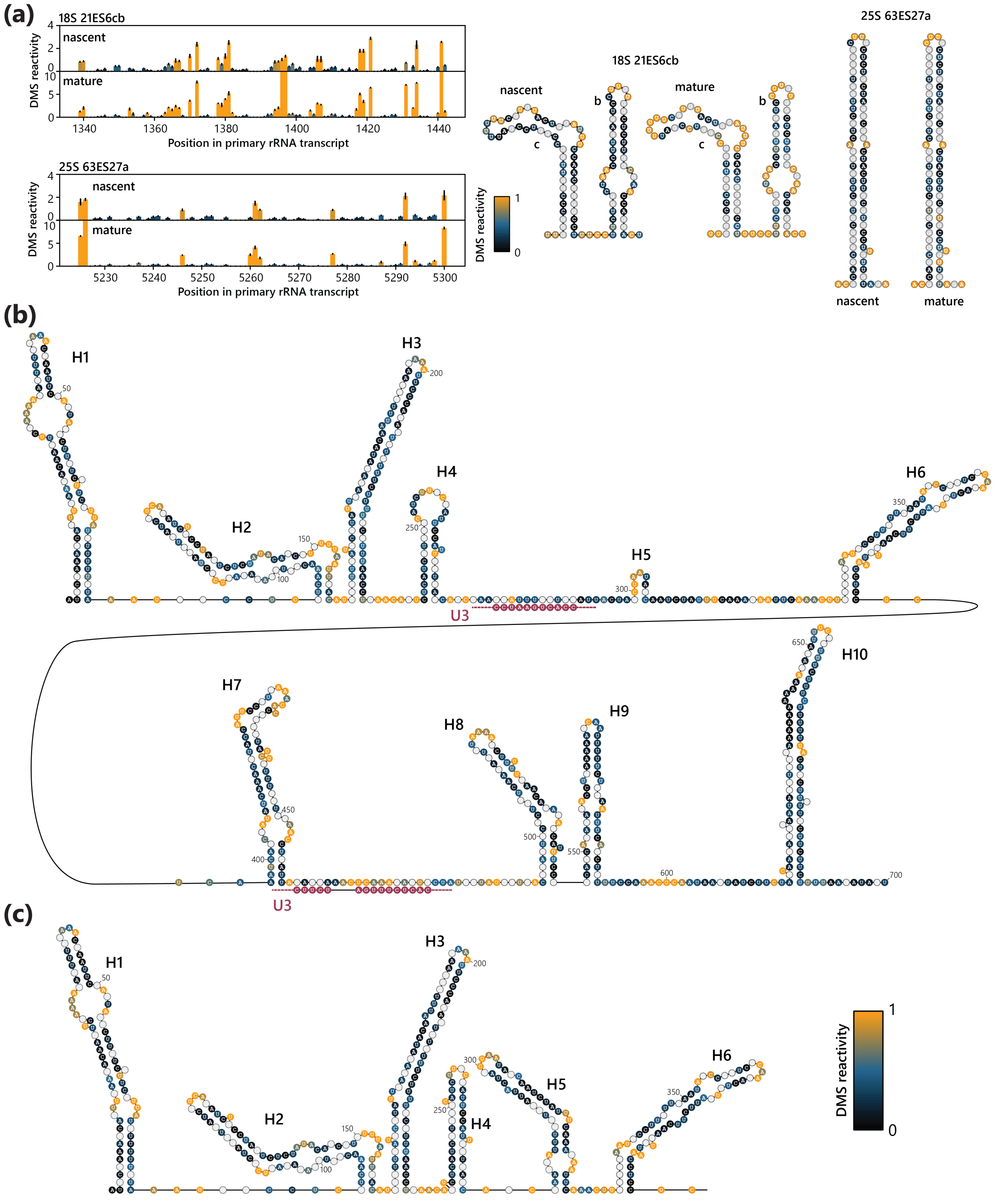


**Fig. S3** *(Related to Fig. 3):* Co-transcriptional formation of rRNA 5'-ETS stem loops. **(a)** DMS reactivity profiles comparing nascent (projected) and mature ribosomal RNA for three stem loops that show high correlation coefficients between nascent and mature states. Error bars show standard deviation of three biological replicates at each position, n=3 for both data sets. The DMS reactivities are overlayed on models of the mature rRNA secondary structure. **(b)** Projected nascent RNA DMS reactivities overlayed on structure models according to cryo-EM structures of 90S pre-ribosomes[31,32]. Discontinuous regions of U3 snoRNA base pairing are indicated but are evidently not base paired in the co-transcriptional window examined (i.e. within ~400 bp of transcription). Wildtype replicate data merged, n=8. **(c)** Stem loops H1-H6 modeled after CoSTseq data without constraints from previous models.

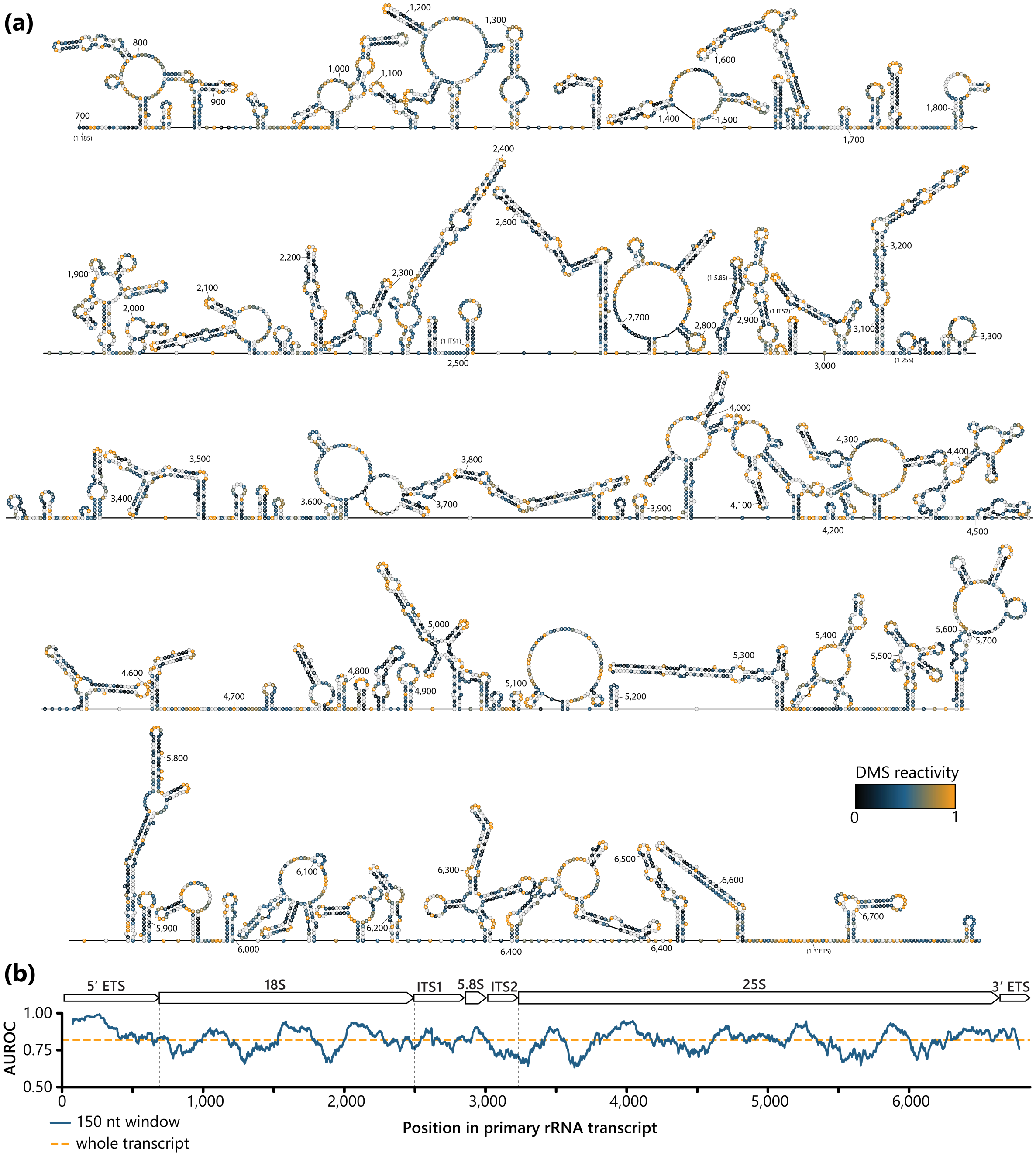


**Fig. S4** *(Related to Fig. 3)***:** Local structure prediction guided by CoSTseq data for the primary rRNA transcript. **(a)** Predicted secondary structure with a maximum base pairing distance of 150 nt. Only wildtype data used, n=8 **(b)** Area under the receiver operating characteristic curve (AUROC) in 150 nt windows to estimate local performance of the predicted model. AUROC for entire transcript: 0.82.

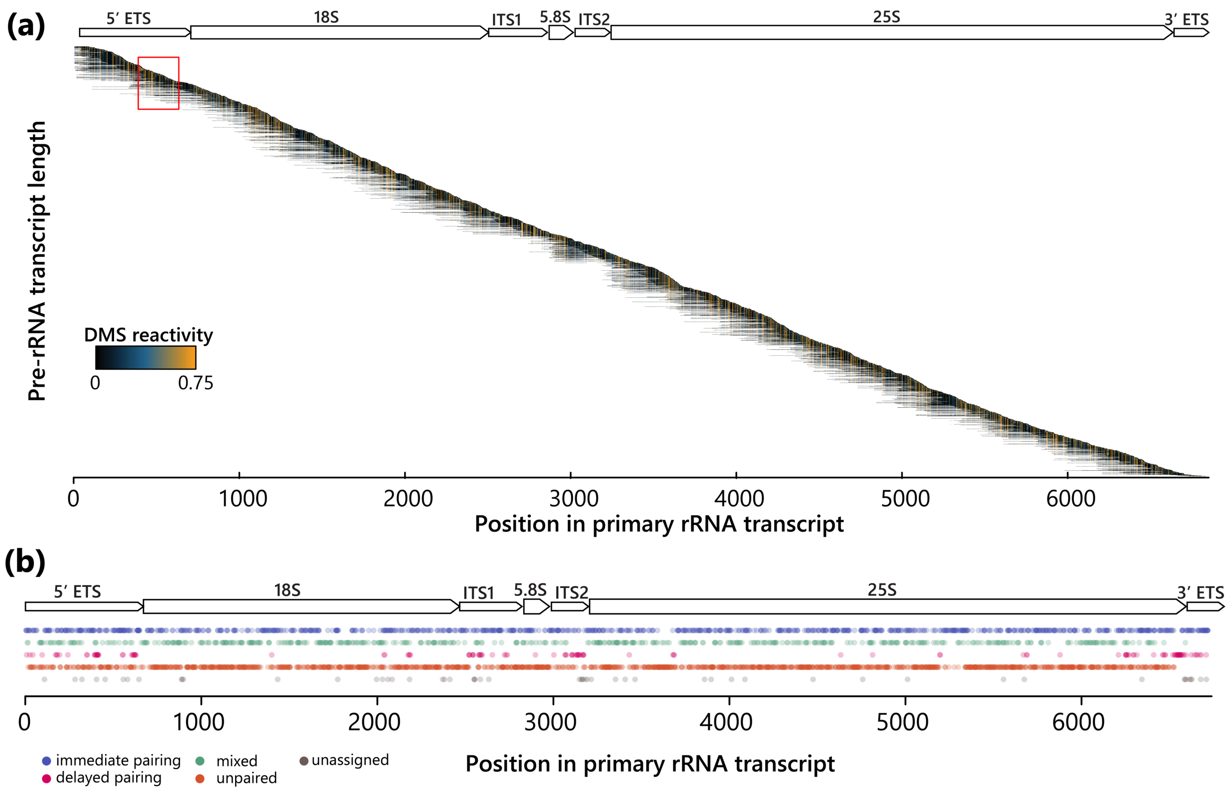


**Fig. S5** *(Related to Fig. 5)***:** Classification of co-transcriptional base pairing. **(a)** Co-transcriptional folding matrix for the entire rDNA gene. Each row represents the DMS reactivity profile of the nascent pre-rRNA at the respective RNA Pol I position within the 5'-ETS, as indicated by the position of the profile along the *y*-axis. Wildtype, n=5. **(b)** Class identity of nucleotides across the pre-rRNA.

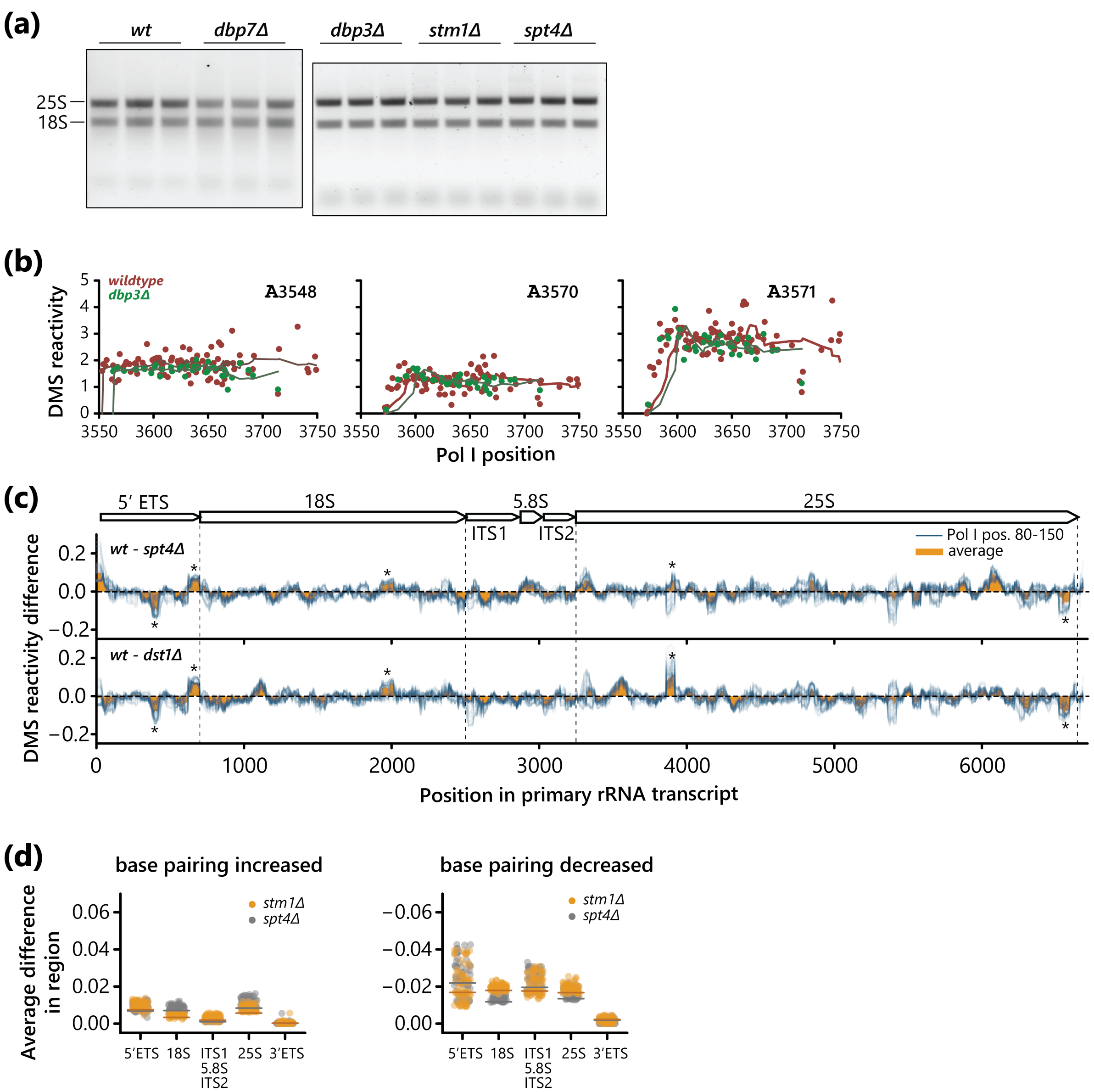


**Fig. S6** *(related to Fig. 6)***:** Co-transcriptional activity of Dbp7 maintains rRNA processing. **(a)** Large and small ribosomal RNA levels for wildtype and mutant strains assessed by agarose gel electrophoresis. **(b)** DMS reactivity trajectories for four nucleotides in wildtype and *dbp3Δ* strains (compare to Fig. 6b). Solid line corresponds to the weighted moving average. **(c)** DMS reactivity difference between wildtype and elongation factor mutants. Each blue line represents data from reads where RNA Pol I was between 80 and 150 nucleotides past the nucleotide of interest, one line for each relative RNA Pol I position. The average difference between all RNA Pol I positions 80-150 is shown in orange. Data for replicates were merged, wildtype: n=5, *spt4Δ*: n=3, *dst1Δ*: n=3. **(d)** Regional average of average (across Pol I positions 80-150) DMS reactivity difference between wildtype and mutants.

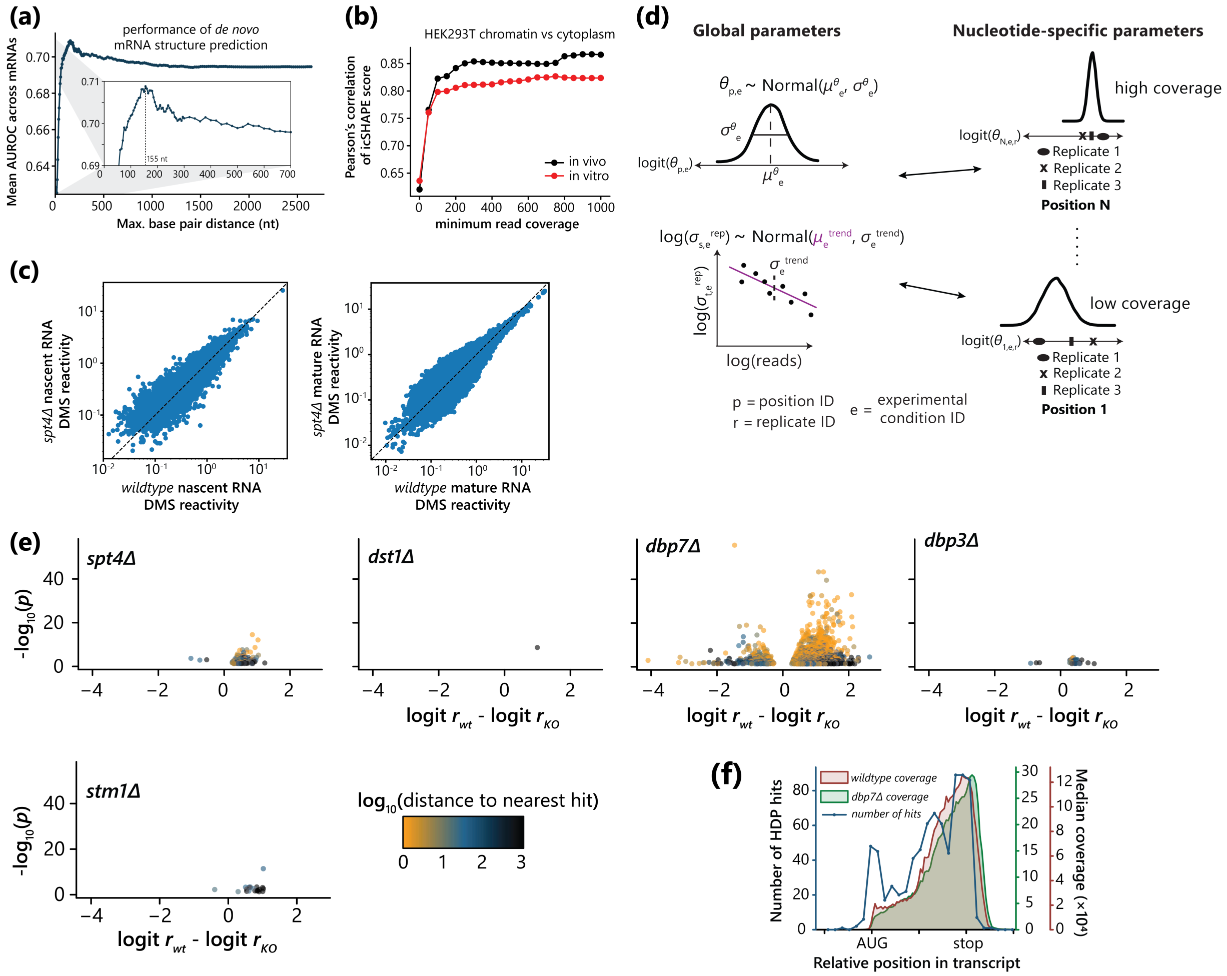


**Fig. S7** *(related to Fig. 7)*: Co-transcriptional activity of Dbp7 maintains rRNA processing. **(a)** *De novo* structure prediction performance assessed by area under the receiver operator characteristic (AUROC) for models predicted with different maximum base pairing distance parameters for all sufficiently covered mRNA transcripts. AUROC is assessed by comparing distributions of DMS reactivities for predicted paired and predicted unpaired nucleotides taken from DMS-MaPseq data. Data from wildtype with three replicates were included. **(b)** Pearson's correlation coefficient for chromatin-associated vs cytoplasmic icSHAPE score, calculated from data by Sun et al. in HEK293T cells as a function of read coverage. **(c)** Scatter plots for DMS reactivities for all sufficiently covered nucleotides from wildtype vs *spt4Δ* strains from CoSTseq and DMS-MaPseq experiments. Replicate data sets were merged, CoSTseq wildtype: n=8, CoSTseq *spt4Δ*: n=3, DMS-MaPseq wildtype: n=3, DMS-MaPseq *spt4Δ*: n=3. **(d)** Principle of HDProbe. Partial pooling of DMS reactivities and replicate variabilities is performed to make use of the high-throughput nature of CoSTseq or DMS-MaPseq data and increase the statistical power of comparisons. **(e)** Volcano plot showing adjusted p-values and mutation rate difference for wildtype vs mutant strains for mRNA (DMS-MaPseq). Data points are colored according to the distance to the nearest significant hit. **(f)** Relative position of significant HDProbe hits within mRNA transcripts in the *dbp7Δ* strain (blue). Median coverage is shown in the same way to assess bias. DMS-MaPseq replicates were combined, wildtype: n=3, *dbp7Δ*: n=3.
